# Context-dependent modulation of social space by dopamine receptors in the *Drosophila melanogaster* mushroom body

**DOI:** 10.64898/2026.05.27.728280

**Authors:** MR Evans, AF Simon

**Author notes:** Correspondence (AFS).

## Abstract

Social behavior takes many forms, yet the fundamental principles of social circuit function are thought to be evolutionarily conserved. Foundational behaviors that precede more complex interactions can reveal the mechanisms underlying these circuits. Social spacing, the regulation of preferred inter-individual distances, is one such behavior and can be quantified in the genetically tractable model *Drosophila melanogaster*. Dopamine and the mushroom body brain region regulate spacing in Drosophila, and extensive dopaminergic signaling occurs within this structure, but receptor-level contributions remain unclear. This report examines how the four dopamine receptors (DopEcR, Dop1R1, Dop1R2, Dop2R) mediate mushroom body-targeted dopaminergic signaling during spacing. Manipulating receptor expression in the entire mushroom body or specific lobes revealed spacing effects that depended on genetic background, receptor identity, sex, and lobe. Receptors could be organized into two opposing pairs: DopEcR and Dop1R1 manipulations increased inter-individual distance, whereas Dop1R2 and Dop2R manipulations decreased inter-individual distance. However, each receptor produced a distinct pattern of effects when male and female data were analyzed across all three mushroom body lobes. These findings support a model in which distributed dopaminergic input engages all four receptors to shape context-dependent spacing decisions, offering insight into social circuit organization in Drosophila and beyond.

**Article Summary:** This study examined how dopamine receptors in the mushroom body (MB) of *Drosophila melanogaster* regulate social spacing, the distance between individuals in groups. The expression of four dopamine receptors was manipulated in different MB lobes to identify their contributions to spacing behavior. The receptors showed distinct effects that depended on receptor type, sex, and MB region. Two receptors increased spacing between flies, while two decreased spacing. These findings support a circuit model in which distributed dopaminergic signals are differentially weighted to generate context-dependent spacing decisions. This work may provide insight into regulation of social spacing in other organisms.

## Introduction

Social behavior can be broadly defined as interactions involving two or more organisms, in which the actions of one individual influence the internal state or subsequent response of another (Sokolowski et al., 2010; Chen & Hong, 2018). These behaviors rely on signaling across neural circuits that continuously integrate social cues with environmental context and internal state, enabling organisms to generate contextually appropriate responses in social settings (Sokolowski et al., 2010; Kim et al., 2017; Chen & Hong, 2018). Disruptions in this process of cue integration can result in behaviors that are expressed at inappropriate times or intensities, which can be detrimental to both the individual and the group (Chen & Hong, 2018).

Although the specific forms and functions of social behavior vary across species, sociality itself is a pervasive feature of life (Sokolowski et al., 2010). To better understand the mechanisms underlying social behavior, it is useful to begin by examining the genetic and neural bases of foundational behaviors that precede and facilitate more complex social interactions.

One such behavior is social spacing, defined as the process by which organisms adjust their position relative to others to maintain an optimal inter-individual distance within a group (Kummer, 1970; Waser & Wiley, 1979; Mogilner et al., 2003; Couzin, 2009; Simon et al., 2012). By governing how individuals distribute themselves in relation to conspecifics, social spacing lays the groundwork for more complex social interactions and serves as a useful model for investigating the neural and genetic mechanisms underlying social behavior. Social spacing is observed across many species (Mogilner et al., 2003). In humans, for example, the regulation of interpersonal distance is an automatic process often described as maintaining a “personal bubble” (Kennedy et al., 2009; Vieira & Marsh, 2014; Lough et al., 2015). Comparable spacing behaviors have been documented in schools of fish, flocks of birds, and even the vinegar fly *Drosophila melanogaster* (Conder, 1949; Emlen, 1952; Breder, 1954; Miller & Stephen, 1966; Simon et al., 2012).

Unlike many other organisms, Drosophila offers an experimentally tractable system in which social spacing can be readily isolated and quantified under controlled conditions (Simon et al., 2012). In laboratory populations of the Canton-Special (CS) strain, individuals consistently maintain separations of roughly two body lengths. The reproducibility of this behavior, combined with the ease of genetic manipulation in Drosophila, makes it a powerful model for investigating the mechanisms underlying social space regulation.

A key neural structure implicated in the regulation of social spacing in Drosophila is the mushroom body (MB) (Burg et al., 2013; Robinson et al., 2024). The MB is a bilaterally symmetrical brain structure best known for its role in learning and memory and is often considered functionally analogous to the mammalian hippocampus (Strausfeld et al., 1998; Sokolowski et al., 2010; Strausfeld & Hirth, 2013; Li et al., 2020). It consists of three major lobes (αβ, α′β′, and γ), each of which can be further subdivided into five discrete compartments that exhibit functional specialization in several contexts (**Supp. Figure 1**) (Aso et al., 2014a; Sitaraman et al., 2015; Hattori et al., 2017; Senapati et al., 2019; Li et al., 2020). MB-targeted manipulations such as neuronal silencing, hyperactivation, and genetic mutants have revealed that disrupted MB function impairs typical social spacing, positioning the MB as a plausible integrative hub in the neural circuitry governing this behavior (Burg et al., 2013; Robinson et al., 2024). In addition to the MB, the neuromodulator dopamine has also been shown to influence social spacing in both males and females, with evidence of sexually dimorphic effects (Fernandez et al., 2017; Yost et al., 2024).

The MB is extensively innervated by dopaminergic neurons (DANs), and dopamine acts as a critical modulator of its neural activity and output (Mao & Davis, 2009; Aso et al., 2014a; Li et al., 2020). Approximately 130 DANs target the MB, and they can be classified into 22 distinct cell types based on their axonal projection patterns (Aso et al., 2014a). Two types originate from the PPL2 cluster and project to the MB calyx. The remaining twenty types arise from the PAM and PPL1 clusters and innervate specific compartments within the MB lobes to modulate synaptic transmission between intrinsic MB Kenyon cells (KCs) and mushroom body output neurons (MBONs) (**Supp. Figure 2**) (Aso et al., 2014a). In learning and memory paradigms, these DANs encode stimulus valence: PAM neurons typically convey reward signals, whereas PPL1 neurons convey punishment. Beyond valence, DAN activity within the MB also reflects a variety of contextual cues, including novelty, expectation, internal physiological state (such as hunger or thirst), locomotor context, and reward prediction error (Krashes et al., 2009; Boto et al., 2014; Hattori et al., 2017; Senapati et al., 2019; Zolin et al., 2021).

The anatomical and functional convergence of the MB and dopaminergic systems, together with their independent links to social spacing, suggests that dopaminergic signaling within the MB may be a critical component of the neural circuitry underlying social space. Dopamine released by DANs modulates KC activity by binding to dopamine receptors expressed on KC membranes (**Supp. Figure 3**) (Aso et al., 2014a). Drosophila encodes four such receptors (DopEcR, Dop1R1, Dop1R2, and Dop2R), which collectively mediate the diverse effects of dopamine throughout the brain (Karam et al., 2020). However, the specific roles of these receptors within the MB during social spacing have not yet been defined, making them strong candidates for uncovering the molecular and circuit-level mechanisms through which dopamine shapes social spacing behavior.

All four dopamine receptors are metabotropic G-protein-coupled receptors (GPCRs), which are membrane proteins that activate G proteins to initiate intracellular signaling cascades upon ligand binding (Yamamoto & Vernier, 2011). Across species, dopamine receptors are divided into two conserved classes, D1 and D2, which are distinguished by their opposing regulation of adenylyl cyclase (AC) and consequent effects on cyclic adenosine monophosphate (cAMP) (**Supp. Figure 4**) (Callier et al., 2003; Yamamoto & Vernier, 2011). In Drosophila, Dop1R1 and Dop1R2 are classified as D1-class receptors because they elevate cAMP levels and promote neuronal excitability upon dopamine binding, though Dop1R2 has a higher activation threshold compared to Dop1R1 (Karam et al., 2020; Driscoll et al., 2021). In contrast, Dop2R is classified as a D2-like receptor because it triggers a decrease in cAMP levels and promotes membrane hyperpolarization (Callier et al., 2003;Yamamoto & Vernier, 2011; Karam et al., 2020). The final receptor, DopEcR, can bind to both dopamine and the steroid hormone 20-hydroxyecdysone (20E) and does not fall within either of the conserved D1 or D2 receptor classes (Karam et al., 2020). Although it shares some properties with D1-class receptors, such as increasing cAMP levels and functioning as an excitatory postsynaptic receptor, its ability to bind multiple ligands and engage diverse intracellular signaling cascades distinguishes it from the other three receptors in Drosophila (Srivastava et al., 2005; Karam et al., 2020).

The four dopamine receptors differ not only in their responses to dopamine, but also in the specific dopaminergic signals they are positioned to detect. Each receptor exhibits a distinct expression profile, with both transcript and protein levels varying across the MB depending on receptor type (Han et al., 1996; Kim et al., 2003; Croset et al., 2018; Kondo et al., 2020). The majority of Kenyon cells (71.5%) express between one and three receptor transcripts in various combinations, with Dop2R expressed at comparatively lower levels than the other receptors (Croset et al., 2018). At the protein level, Dop1R1 and Dop2R are broadly distributed across Kenyon cell membranes (**Supp. Figure 5**) (Kondo et al., 2020). In contrast, DopEcR is enriched at the axon initial segment near the calyx, whereas Dop1R2 localizes to complementary anterior peduncle and distal axonal regions. These distinct expression patterns coupled with downstream signaling differences likely underlie receptor-specific roles in behavior, many of which have been well characterized (**Supp. Table 1**; Karam et al., 2020). As a result, each receptor represents a distinct candidate when studying dopaminergic modulation.

To investigate the role of individual dopamine receptors in Drosophila social spacing, RNA interference (RNAi) was used to selectively manipulate receptor expression across the mushroom body (MB), both globally and within specific lobes. These manipulations revealed two functional groupings of receptors based on their effects on spacing behavior: when effects were observed, altering DopEcR or Dop1R1 expression caused flies to settle farther apart, whereas altering Dop1R2 or Dop2R expression resulted in flies moving closer together. Although all four receptors contributed to social spacing, the pattern of effects depended on receptor identity, sex, and the MB region targeted. Based on these findings and the known architecture of MB circuits, a model is proposed to account for the widespread and sexually dimorphic contributions of dopamine receptors to social spacing. Given the evolutionary conservation of dopaminergic systems and social behavior pathways, these findings may provide insight into the regulation of social spacing in other organisms (Reaume & Sokolowski, 2011; Yamamoto & Vernier, 2011; Goodson & Kingsbury, 2013).

## Materials and Methods

### Parental Fly Lines

All driver, effector, and control lines used in genetic crosses were obtained from the Bloomington Drosophila Stock Center (BDSC; Department of Biology, Indiana University, Bloomington, IN, NIH P40OD018537). All driver lines targeting the mushroom body (MB) region were contributed to BDSC by the same donor (Gerald M. Rubin, Howard Hughes Medical Institute, Janelia Research Campus) and share a common genetic background, hereafter referred to as the Janelia background. Effector lines carrying UAS-linked RNAi constructs targeting each dopamine receptor were originally generated by the Transgenic RNAi project (TriP) at Harvard Medical School. The corresponding control line contains an empty attP2 docking site and shares the same genetic background as the four TRiP lines, hereafter referred to as the TRiP background. A GFP effector line was also used to confirm driver expression profiles. Details of all parental lines are provided in **Supp. Table 2**.

### Receptor Manipulation

Experimental lines with altered receptor expression were generated by crossing MB– and lobe-specific driver lines to RNAi effector lines. Corresponding genetic background controls were generated by crossing driver lines to the TRiP control line. For all crosses, 20 virgin females from one parental fly line and 20 males from the other were combined in a bottle and allowed to mate for 7 days. Parental flies were anesthetized with carbon dioxide to facilitate collection and were no older than 14 days at the time of crossing to control for behavioral variation in progeny of aged flies (Brenman-Suttner et al., 2018). Adults were removed prior to progeny eclosion. Genetic crosses used to generate experimental and control lines are summarized in **Supp. Table 3**, with crossing schemes further illustrated in **Supp. Figure 6**.

### Rearing Conditions

All Drosophila lines and crosses were maintained in mixed-sex bottles containing handmade food prepared following the Fisher Scientific Jazz-Mix^TM^ recipe [brown sugar (118.13 g/L), corn meal (30.24 g/L), yeast (17.48 g/L), agar (5.67 g/L), benzoic acid (1.89g/L), methyl paraben (0.71 g/L), propionic acid (0.71 g/L)]. Flies were reared in an incubator within the insect module of the Biotron Facility at Western University under controlled conditions of 25°C, 50% relative humidity, and a 12:12 h light:dark cycle (lights on at 9:00am and off at 9:00pm).

### Testing Efficacy of Driver Expression

To confirm targeted expression, each driver was crossed to a *GFP* effector line carrying a transgene encoding Green Fluorescent Protein (GFP) fused to the mouse-derived membrane protein mDC8. The resulting *driver>GFP* progeny express membrane-localized GFP under the control of the driver, allowing for visualization of the driver’s expression pattern by fluorescence microscopy. Whole brains of 3-4-day old *driver>GFP* flies were dissected in 1x phosphate-buffered saline (PBS) and mounted on glass slides using FluoroShield^TM^ mounting media (Sigma-Aldrich, catalog number NC0375254). *GFP* and *attP2* flies were prepared in parallel as negative controls, as neither fly line is expected to express GFP. For each genotype, five male and five female brains were dissected. Imaging was performed on an Imager Z1 Zeiss compound fluorescent microscope at 10x or 20x magnification, using excitation and emission wavelengths of 488nm and 507nm respectively. Although Drosophila brain lipids display background autofluorescence, GFP expression is distinguishable as brighter green fluorescence above this background. Representative brain images for each driver are found in **Supp. Figure 7**.

### Fly Handling Prior to Behavioral Assays

Progeny from each cross were kept in mixed-sex bottles with food to allow mating prior to behavioral testing, as virginity is an unnatural physiological state known to influence behavior (Simon et al., 2012). Twenty-four hours before testing, 2-3-day old flies were briefly cold-anesthetized and sorted into same-sex groups of 12-18 individuals in preparation for the social space assay. Group sizes were optimized in previously established protocols, and flies were separated by sex to enable between-sex comparisons (McNeil et al., 2015; Simon et al., 2012). Sorted groups were transferred to vials of fresh food and left overnight to recover from any residual effects of anesthesia. On the morning of testing, flies were again transferred to fresh food and given 2 hours to acclimate to experimental conditions (25°C, 50% relative humidity). All social space assays were performed in a dedicated behavior room under uniform lighting to standardize visual conditions, with testing restricted to 12:00pm-4:00pm to minimize behavioral variability associated with circadian rhythm.

### The Social Space Assay

The social space assay was performed as previously described in Simon et al. (2012) and McNeil et al. (2015). Groups of 12-18 flies, 3-4-days old and of the same genotype and sex, were gently funneled into a vertically oriented two-dimensional triangular chamber. The chamber was tapped three times on the lab bench to bring all flies to the bottom, establishing a uniform starting position. This tapping also elicited negative geotaxis, an innate escape response in which flies climb upward against gravity after disturbance (Gargano et al., 2005). As flies ascend, the triangular shape of the chamber forces them into close proximity at the top, prompting individuals to spread out and establish their preferred spacing. Flies were allowed to move freely until the group stabilized, which typically occurred after 20-40 minutes, at which point pictures were taken for analysis. Captured images were analyzed with the open access software ImageJ (RRID:SCR_003070 – Schneider et al., 2012) using plug-ins published in Yost et al. (2020) and available at https://github.com/flugrugger/bubble (BubbleMacroWalkthrough.docx, Bubble.txt, SocialSpaceAnalysisonImageJ-2019.doc). Social space was quantified by counting the number of flies located within a four-body-length (4BL) radius of each individual in the image and calculating the mean value across all flies in the chamber. Each chamber containing 12-18 flies constituted one replicate. Image analysis data were imported to GraphPad Prism 10 software for statistical analysis and graphical representation.

### Statistical Analysis

Data from behavioral assays were graphed and analyzed using GraphPad Prism (RRID:SCR_002798, version 10.5.0 for Mac, GraphPad Software, La Jolla California USA, www.graphpad.com). Outliers were identified and removed using the ROUT method (Robust regression and Outlier removal; Motulsky & Brown, 2006). Normality was assessed for all genotype × sex groups within each graph prior to applying parametric tests. A distribution was considered approximately Gaussian if it passed at least half of the following four normality tests: D’Agostino-Pearson omnibus, Anderson-Darling, Shapiro-Wilk, and Kolmogorov-Smirnov. Two-way ANOVAs were used to evaluate the effects of both genotype and sex on the dependent variable (flies/4BL) as well as potential interactions between these factors. Main effects and interaction effects from the two-way ANOVAs are displayed on the corresponding graphs, with p-values below the alpha threshold of 0.05 bolded to indicate clear statistical evidence of an effect. Šidák’s post-hoc tests were performed to compare experimental and control genotypes within the same sex. P-values from post-hoc tests are shown on graphs only when relevant to interpreting the main and interaction effects. Full details of all statistical analyses, including test statistics and p-values, are provided in **Supp. Table 4**.

### Social Space Effect Size Quantification

Effect sizes were calculated separately for males and females within each experimental dataset. For each driver, the experimental effect size was defined as the difference in the mean number of flies within four body lengths (flies/4BL) between experimental and control genotypes (*driver>RNAi* − *driver/attP2*), where negative values indicate increased social spacing and positive values indicate decreased spacing. To account for baseline effects of RNAi constructs, baseline effect sizes were calculated in the same manner from a control dataset (*RNAi* − *attP2*). Additive effect sizes were obtained by subtracting the baseline effect from the experimental effect, isolating the driver-dependent component of behavioral change (**Supp. Figure 8**).

Additive effect sizes were normalized by dividing by the mean flies/4BL of the corresponding experimental control genotype (*driver/attP2*), yielding a normalized percent change in social spacing to facilitate comparisons across datasets. In one case where the baseline effect was much larger in magnitude than the experimental effect, the normalized value exceeded 100%; this value reflects the mathematical adjustment and was retained for completeness, though emphasis should be placed on direction and relative magnitude of change rather than absolute percentage. All effect sizes and normalized values are provided in **Supp. Table 5**.

### Drawings

Made using BioRender University of Western Ontario site license.

## Results

All experimental social space datasets (*MB, αβ, α′β′, γ*) are presented alongside the control dataset comparing baseline behavior of the *DopR-RNAi* lines to the *attP2* control. In some cases, the presence of the RNAi construct produced behavioral effects even in the absence of a driver, likely due to construct insertion or leaky expression. To ensure accurate interpretation, all experimental findings are evaluated relative to baseline effects in the control dataset to account for any pre-existing behavioral perturbations.

Across all genotypes, females displayed closer social spacing than males (two-way ANOVA: effect of sex, p < 0.05 for all genotypes, apart for *αβ>Dop2R-RNAi* p=0.0600). This sex effect is consistent with previous reports (Robinson et al., 2024). Since this difference was robust and expected, the following text focusses on genotype and interaction effects.

### DopEcR manipulation in the MB may increase social space in males only

When *DopEcR* expression was manipulated in the MB within the Janelia background, both sexes showed an apparent increase in flies/4BL relative to controls, indicating flies were settling closer together (two-way ANOVA: effect of genotype, *p = 0.0114*; effect of genotype × sex interaction, *p = 0.2480*; **Figure 1B**). However, in the control dataset, *DopEcR-RNAi* flies were also closer than *attP2* controls at baseline (two-way ANOVA: effect of genotype, *p* < 0.0001; effect of genotype × sex interaction, *p = 0.2384*; **Figure 1A**), indicating the RNAi construct influences behavior even in the absence of a driver.

**Figure 1.**
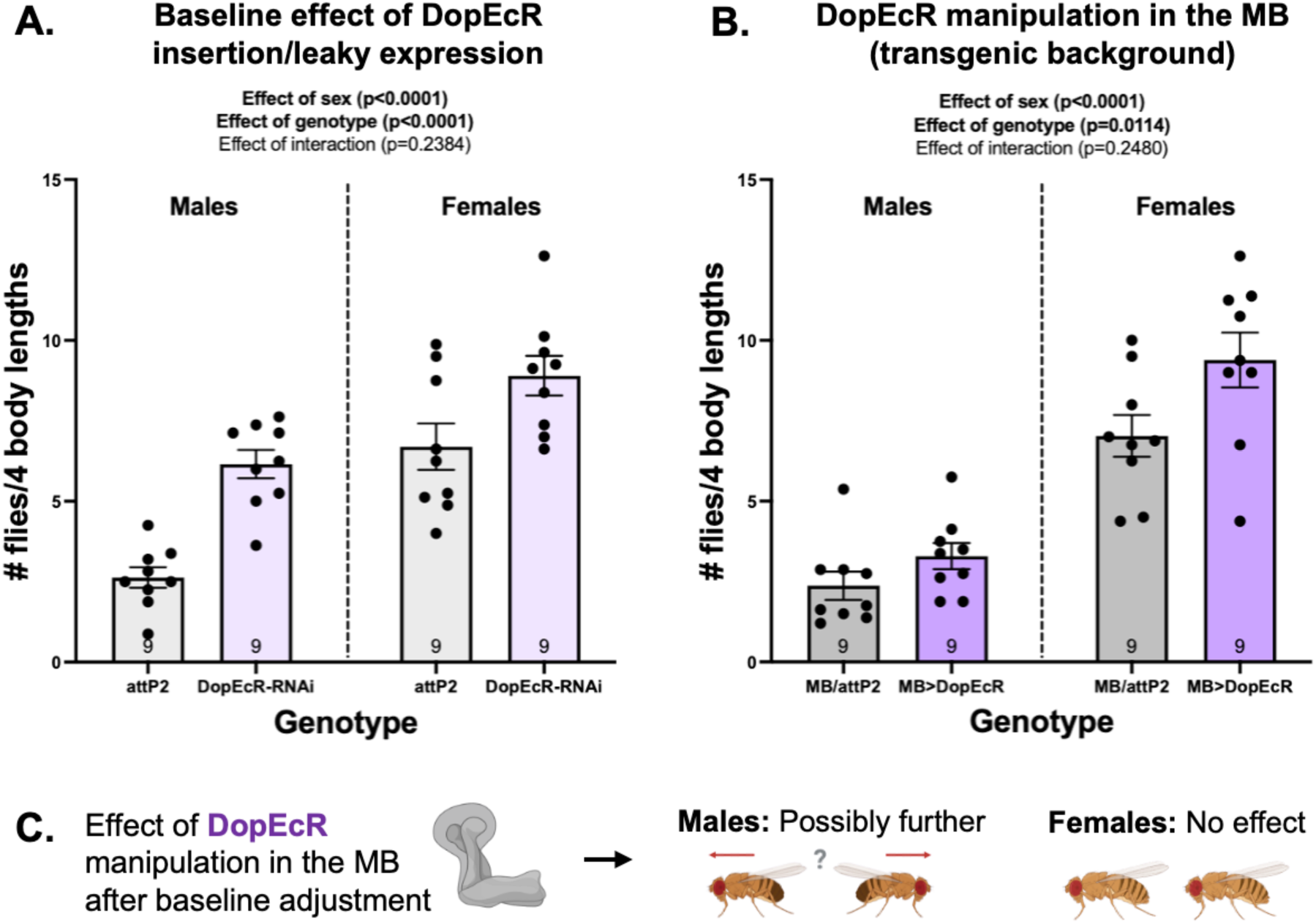
*DopEcR* manipulation in the MB may increase male social space. Bars show the mean number of flies within four body lengths (± SEM) for each genotype, with lower values indicating farther/increased spacing. Each dot represents one chamber of 12-18 flies (N=9 chambers per genotype). Data were analyzed by two-way ANOVA (factors: sex and genotype). **A.** Control dataset comparing *DopEcR-RNAi* flies with *attP2* controls to assess potential baseline effects of construct insertion or leaky expression. *DopEcR-RNAi* flies were closer in both sexes (effect of genotype, *p < 0.0001*). **B.** Experimental dataset depicting *MB>DopEcR-RNAi* within the Janelia background. Prior to baseline correction, *DopEcR* manipulation appeared to decrease spacing in both sexes (effect of genotype, *p = 0.0114*). **C.** After accounting for baseline effects, the data indicate that *MB>DopEcR-RNAi* males may settle farther apart, whereas *MB>DopEcR-RNAi* females show no change in spacing.

In males, both the control and experimental datasets individually showed increased proximity. However, the baseline increase in the control dataset was substantially larger. After adjusting for baseline differences and calculating the additive effect size, the outcome was reversed: male flies with *DopEcR* manipulated in the MB were actually farther apart. This corresponded to a normalized decrease of 110% in flies/4BL. Although this value is not biologically literal, it reflects the mathematical adjustment for baseline effects and is interpreted qualitatively to convey the magnitude of the change produced by *DopEcR-RNAi* in the MB. Despite this large effect size, the conclusion emerging after baseline correction is opposite to what would be inferred from the experimental dataset alone and should therefore be interpreted with caution. The available data indicate that males were possibly farther apart with *DopEcR* manipulation in the MB, but this tentative conclusion should be confirmed with further experiments.

In females, the closer spacing observed in the experimental dataset mirrored the baseline pattern seen in the control dataset, resulting in no overall effect of modifying *DopEcR* expression in the MB after baseline correction.

### Dop1R1 manipulation in the MB may increase social space in males only

When *Dop1R1* expression was manipulated in the MB within the Janelia background, social spacing did not change in either sex in the experimental dataset (two-way ANOVA: effect of genotype, *p = 0.9936*; effect of genotype × sex interaction, *p = 0.6468*; **Figure 2B**). In the control dataset, *Dop1R1-RNAi* flies were also not statistically different than *attP2* flies, though the genotype effect approached significance (two-way ANOVA: effect of genotype, *p = 0.0693*; effect of genotype × sex interaction, *p = 0.2942*; **Figure 2A**). Post-hoc Šidák’s tests indicated a trend toward closer spacing in male *Dop1R1-RNAi* flies relative to controls (Šidák’s test: *p = 0.0886*), suggesting a possible baseline effect of the RNAi construct even in the absence of a driver.

**Figure 2.**
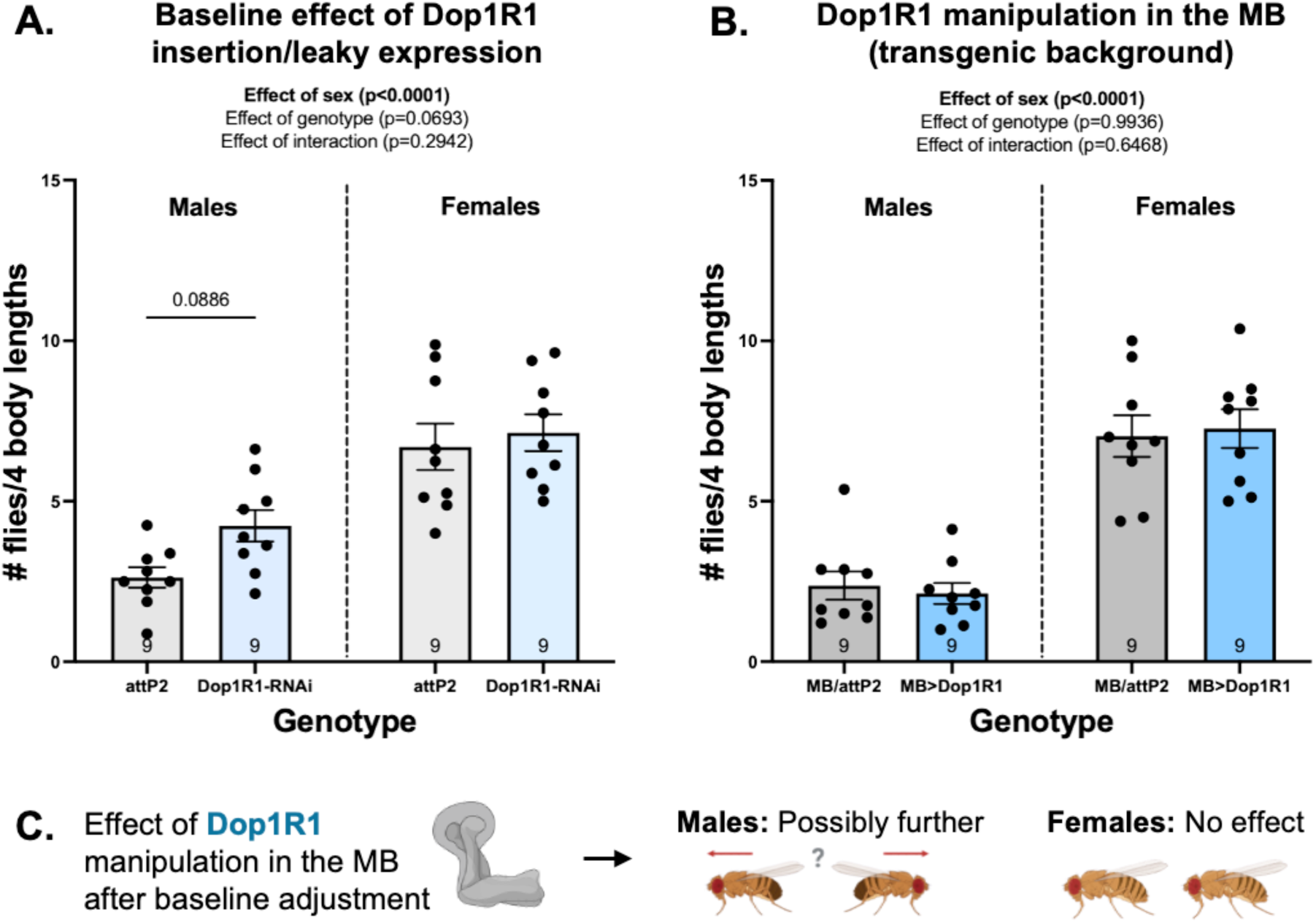
*Dop1R1* manipulation in the MB may increase male social space. Bars show the mean number of flies within four body lengths (± SEM) for each genotype, with lower values indicating farther/increased spacing. Each dot represents one chamber of 12-18 flies (N=9 chambers per genotype). Data were analyzed by two-way ANOVA (factors: sex and genotype). **A.** Control dataset comparing *Dop1R1-RNAi* flies with *attP2* controls to assess potential baseline effects of construct insertion or leaky expression. *Dop1R1-RNAi* flies were not statistically different from controls, though the genotype effect approached significance (effect of genotype, *p = 0.0693*). Post-hoc Šidák’s tests suggest a trend toward closer spacing in males at baseline (*p = 0.0886*). **B.** Experimental dataset depicting *MB>Dop1R1-RNAi* within the Janelia background. Prior to baseline correction, *Dop1R1* manipulation did not appear to alter social space in either sex (effect of genotype, *p = 0.9936*). **C.** After accounting for baseline effects, the data indicate that *MB>Dop1R1-RNAi* males may settle farther apart, whereas *MB>Dop1R1-RNAi* females show no change in spacing.

After accounting for baseline effects, males displayed a normalized percent decrease of 78% in flies/4BL when *Dop1R1-RNAi* was driven in the MB, suggesting that flies settled farther apart with this manipulation. However, this effect became evident only after baseline correction and was not apparent from the experimental dataset alone, warranting caution in its interpretation. The available data indicate that males were possibly farther apart with *Dop1R1* manipulation in the MB, but this tentative conclusion should be confirmed with further experiments.

In females, altering the expression of *Dop1R1* in the MB did not affect social spacing. This conclusion was reinforced by comparison to the control dataset, which showed no baseline effect of the *Dop1R1-RNAi* construct.

### Dop1R2 manipulation in the MB decreases social space in both sexes

When *Dop1R2* expression was manipulated in the MB within the Janelia background, both sexes showed an increase in flies/4BL relative to the control genotype, indicating that flies were settling closer together (two-way ANOVA: effect of genotype, *p* = *0.0206*; effect of genotype × sex interaction, *p = 0.9825*; **Figure 3B**). In the control dataset, *Dop1R2-RNAi* flies were not statistically different than *attP2* flies, though the genotype effect approached significance (two-way ANOVA: effect of genotype, *p = 0.0975*; effect of genotype × sex interaction, *p = 0.2522*; **Figure 3A**). Post-hoc Šidák’s tests indicated a trend toward farther spacing in female *Dop1R2-RNAi* flies relative to controls (Šidák’s test: *p = 0.0986*), suggesting a possible baseline effect of the RNAi construct even in the absence of a driver.

**Figure 3.**
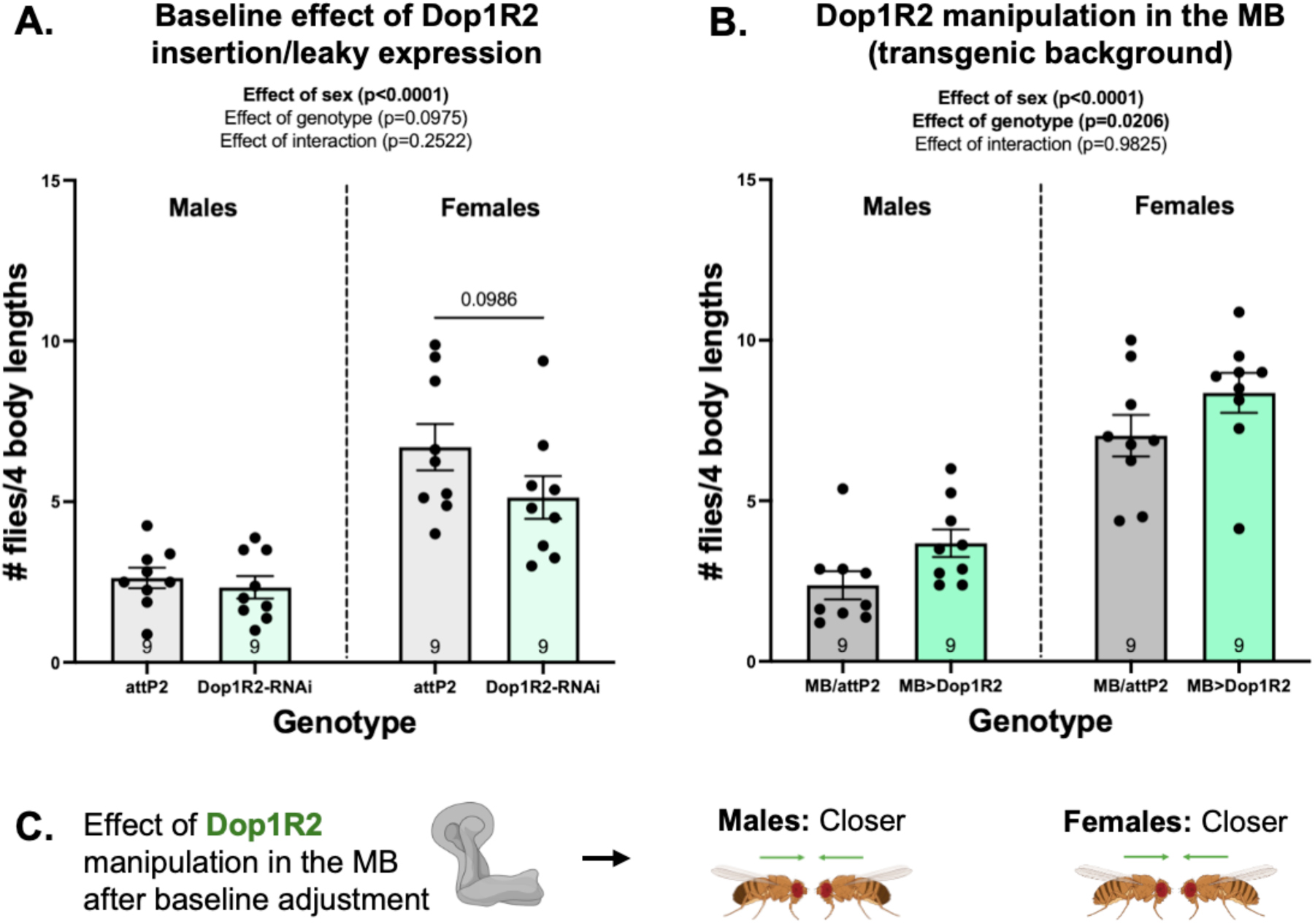
*Dop1R2* manipulation in the MB decreases social space in both sexes. Bars show the mean number of flies within four body lengths (± SEM) for each genotype, with lower values indicating farther/increased spacing. Each dot represents one chamber of 12-18 flies (N=9 chambers per genotype). Data were analyzed by two-way ANOVA (factors: sex and genotype). **A.** Control dataset comparing *Dop1R2-RNAi* flies with *attP2* controls to assess potential baseline effects of construct insertion or leaky expression. *Dop1R2-RNAi* flies were not statistically different from controls, though the genotype effect approached significance (effect of genotype, *p = 0.0975*). Post-hoc Šidák’s tests suggest a trend toward farther spacing in females at baseline (*p = 0.0986*). **B.** Experimental dataset depicting *MB>Dop1R2-RNAi* within the Janelia background. Prior to baseline correction, *Dop1R2* manipulation appeared to decrease spacing in both sexes (effect of genotype, *p = 0.0206*). **C.** After accounting for baseline effects, the data indicate that *MB>Dop1R2* flies of both sexes settle closer together.

After accounting for baseline effects, both sexes exhibited closer social spacing following *Dop1R2* manipulation in the MB, with normalized percent increases in flies/4BL of 68% in males and 41% in females. These effects were consistent with observations from the experimental dataset alone and were reinforced by comparison to the control dataset.

### Dop2R manipulation in the MB decreases social space in both sexes

When *Dop2R* expression was manipulated in the MB within the Janelia background, there was no clear evidence of a change in flies/4BL relative to the control genotype, although both sexes showed a trend toward closer spacing (two-way ANOVA: effect of genotype, *p* = *0.1378*; effect of genotype × sex interaction, *p = 0.8828*; **Figure 4B**). Analysis of the control dataset indicated that the RNAi construct produced a baseline increase in spacing among females but not males (two-way ANOVA: effect of genotype, *p* = *0.0009*; effect of genotype × sex interaction, *p = 0.0325*; **Figure 4A**). Post-hoc Šidak’s tests provided further evidence that female *Dop2R-RNAi* flies were farther apart than *attP2* controls at baseline (Šidak’s test: *p = 0.0005*), even in the absence of a driver.

**Figure 4.**
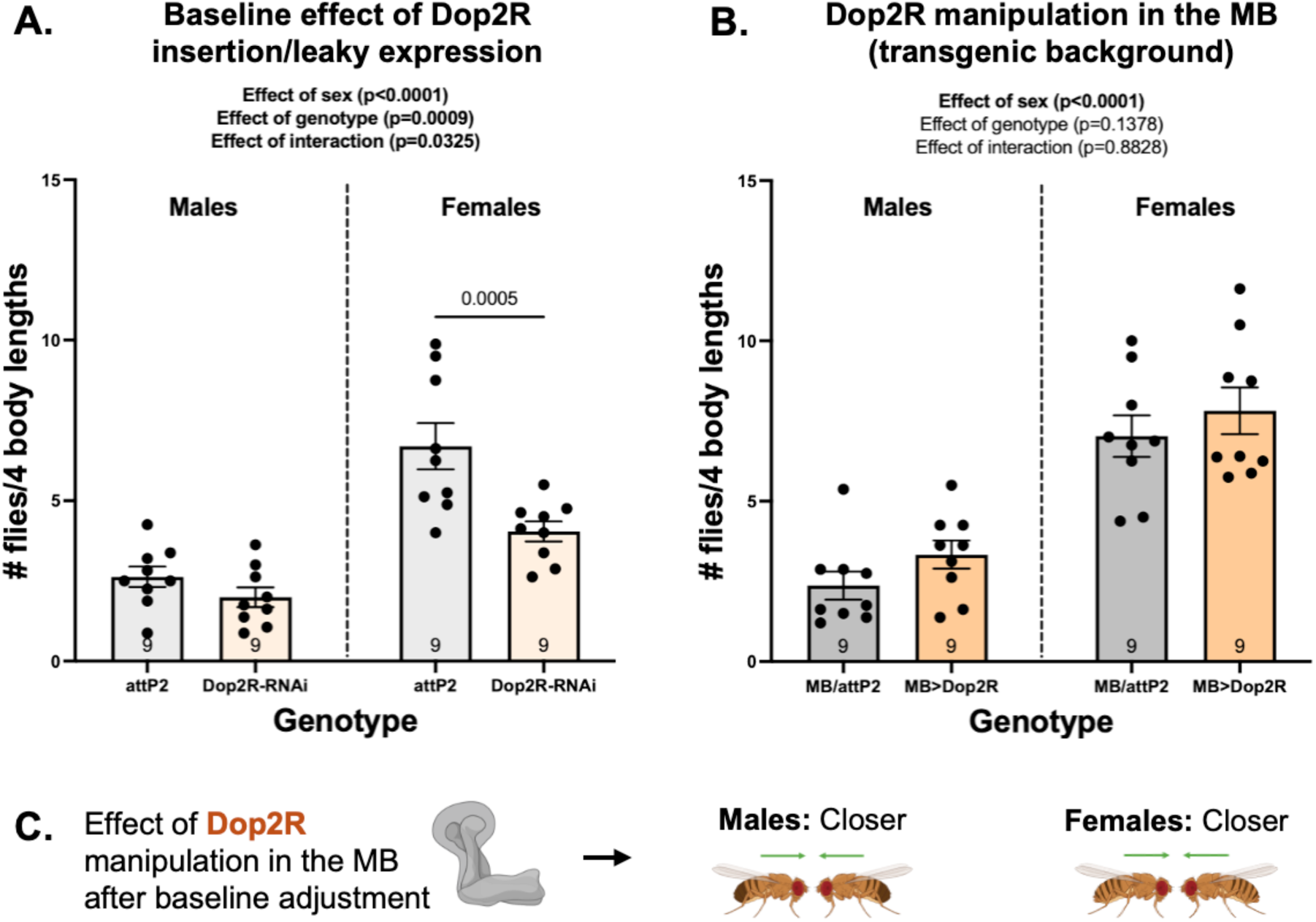
*Dop2R* manipulation in the MB decreases social space in both sexes. Bars show the mean number of flies within four body lengths (± SEM) for each genotype, with lower values indicating farther/increased spacing. Each dot represents one chamber of 12-18 flies (N=9 chambers per genotype). Data were analyzed by two-way ANOVA (factors: sex and genotype). **A.** Control dataset comparing *Dop2R-RNAi* flies with *attP2* controls to assess potential baseline effects of construct insertion or leaky expression. *Dop2R-RNAi* females were farther, while males showed no clear change in spacing (effect of genotype, *p = 0.0009*; effect of genotype × sex interaction, *p = 0.0325*). Post-hoc Šidák’s tests confirmed this baseline effect in females (*p = 0.0005*). **B.** Experimental dataset depicting *MB>Dop2R-RNAi* within the Janelia background. Prior to baseline correction, *Dop2R* manipulation did not appear to alter spacing in either sex, though there was a trend toward closer spacing (effect of genotype, *p = 0.1378*). **C.** After accounting for baseline effects, the data indicate that *MB>Dop2R-RNAi* flies of both sexes settle closer together.

After accounting for baseline effects, both sexes exhibited closer social spacing following *Dop2R* manipulation in the MB, with normalized percent increases in flies/4BL of 68% in males and 49% in females. These effects were consistent with trends observed in the experimental dataset alone and were reinforced by comparison to the control dataset.

### DopEcR manipulation increases female spacing in α′β′ and may increase male spacing in αβ and γ lobes

#### Baseline effect of *DopEcR-RNAi* construct

*DopEcR-RNAi* flies of both sexes settle closer together relative to *attP2* controls, reflecting a baseline effect of the RNAi construct even in the absence of a Gal4 driver (see Section 4.1.1 for details; **Figure 5A**).

**Figure 5.**
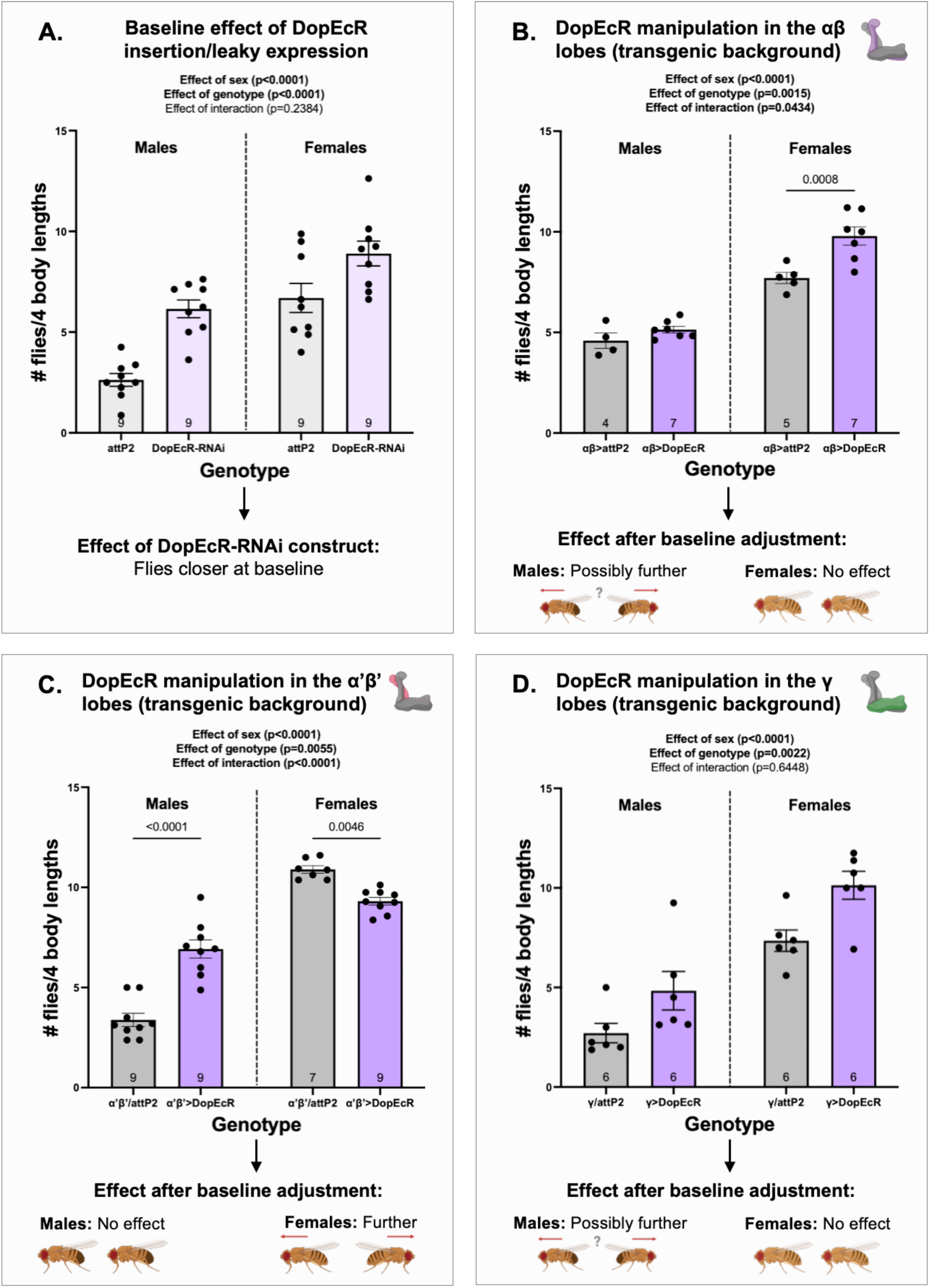
*DopEcR* manipulation increases spacing in lobe– and sex-specific ways. Bars show the mean number of flies within four body lengths (± SEM) for each genotype, with lower values indicating farther/increased spacing. Each dot represents one chamber of 12-18 flies (N=4-9 chambers per genotype). Data were analyzed by two-way ANOVA (factors: sex and genotype); graphs display all main and interaction effects, with post-hoc Šidak’s p-values included when relevant for interpretation. Comparison of *DopEcR-RNAi* flies with *attP2* controls in the control dataset **(A)** reveals a baseline effect of the RNAi construct, with flies settling closer together even in the absence of a Gal4 driver. Manipulating *DopEcR* expression within the αβ **(B)**, α’β’ **(C)**, and γ **(D)** lobes reveals lobe-specific contributions to social spacing within each sex after baseline correction. *DopEcR* may increase social spacing in males when manipulated in the αβ and γ lobes, whereas manipulation in the α’β’ lobes increases spacing in females.

#### αβ lobes

Manipulating *DopEcR* expression in the αβ lobes within the Janelia background decreased social space in females but not males (two-way ANOVA: effect of genotype, *p* = *0.0015*; effect of genotype × sex interaction, *p = 0.0434*; **Figure 5B**). Post-hoc Šidak’s tests provided further evidence that female *αβ>DopEcR* flies were closer than controls (Šidak’s test: *p = 0.0008*). However, this decrease in spacing mirrored the baseline pattern in the control dataset, resulting in no overall effect of *αβ>DopEcR-RNAi* on female spacing after baseline correction. In males, accounting for baseline effects suggested that flies settle farther apart following *DopEcR* manipulation in the αβ lobes, corresponding to a 65% normalized percent decrease in flies/4BL. Since this effect only emerged after baseline correction, it is tentatively concluded that *DopEcR* manipulation in the αβ lobes may increase social spacing in males.

#### α’β’ lobes

Manipulating *DopEcR* expression in the α’β’ lobes within the Janelia background produced opposite effects across sexes: males exhibited decreased social spacing, whereas females showed increased spacing (two-way ANOVA: effect of genotype, *p* = *0.0055*; effect of genotype × sex interaction, *p < 0.0001*; **Figure 5C**). Post-hoc Šidak’s tests supported these effects in both males (*p < 0.0001*) and females (*p = 0.0046*). However, the decreased spacing observed in males mirrored the baseline pattern in the control dataset, resulting in no overall effect of α’β’-lobe *DopEcR* manipulation after baseline correction. In females, comparison to the control dataset reinforced the conclusion that flies settle farther apart following α’β’-lobe *DopEcR* manipulation, with a normalized 35% decrease in flies/4BL after baseline correction.

#### γ lobes

Manipulating *DopEcR* expression in the γ lobes within the Janelia background produced an apparent decrease in social spacing, with both sexes settling closer together relative to controls (two-way ANOVA: effect of genotype, *p* = *0.0022*; effect of genotype × sex interaction, *p = 0.6448*; **Figure 5D**). However, the comparatively larger baseline decrease in spacing caused by the *DopEcR-RNAi* construct reversed the effect in males after baseline correction: rather than settling closer, γ-lobe *DopEcR* manipulation caused males to be farther apart, with a normalized 52% decrease in flies/4BL. Since this effect only emerged after baseline correction, it is tentatively concluded that *DopEcR* manipulation in the γ lobes may increase social spacing in males. In females, the decreased spacing observed in the experimental dataset mirrored the baseline pattern in the control dataset, resulting in no overall effect of γ-lobe *DopEcR* manipulation on female spacing after baseline correction.

#### Summary

In males, *DopEcR* manipulation in both the αβ and γ lobes may cause flies to settle farther apart relative to controls, recapitulating the possible increase in social spacing observed at the whole-MB level. In females, manipulating *DopEcR* expression in the α’β’ lobes increased spacing relative to controls, which was unexpected given that *DopEcR* showed no influence on social space behavior at the whole-MB level.

### Dop1R1 manipulation increases female spacing in α′β′ and may increase male spacing in all lobes

#### Baseline effect of *Dop1R1-RNAi* construct

*Dop1R1-RNAi* females did not differ in spacing relative to *attP2* controls. In contrast, males settled closer together, indicating a male-specific baseline effect of the RNAi construct even in the absence of a Gal4 driver (see Section 4.1.2 for details; **Figure 6A**).

**Figure 6.**
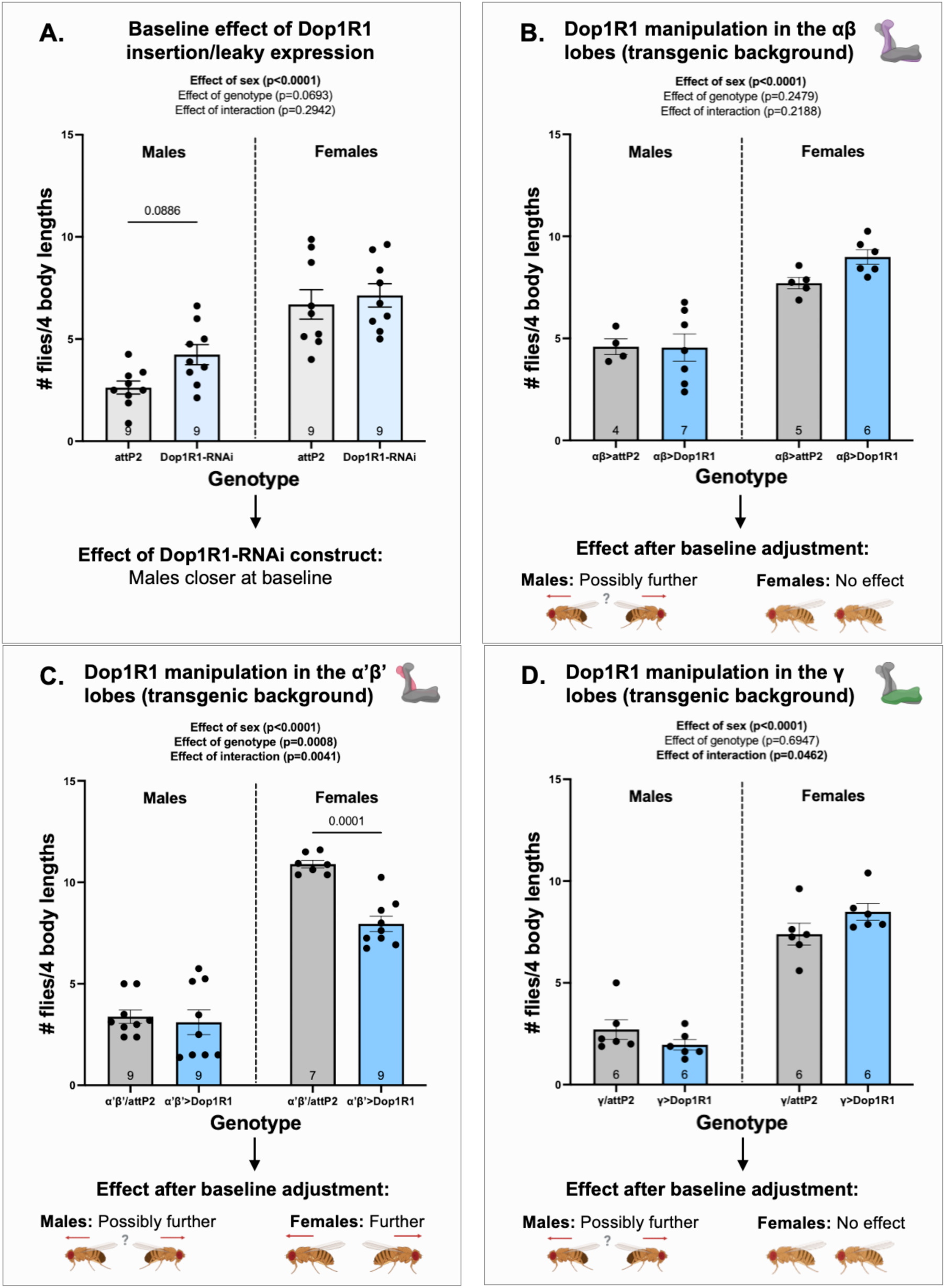
*Dop1R1* manipulation increases spacing in lobe– and sex-specific ways. Bars show the mean number of flies within four body lengths (± SEM) for each genotype, with lower values indicating farther/increased spacing. Each dot represents one chamber of 12-18 flies (N=4-9 chambers per genotype). Data were analyzed by two-way ANOVA (factors: sex and genotype); graphs display all main and interaction effects, with post-hoc Šidak’s p-values included when relevant for interpretation. Comparison of *Dop1R1-RNAi* flies with *attP2* controls in the control dataset **(A)** reveals a baseline effect of the RNAi construct, with males settling closer together even in the absence of a Gal4 driver. Manipulating *Dop1R1* expression within the αβ **(B)**, α’β’ **(C)**, and γ **(D)** lobes reveals lobe-specific contributions to social spacing within each sex after baseline correction. *Dop1R1* may increase social spacing in males when manipulated in any of the three lobes, whereas manipulation in the α’β’ lobes increases spacing in females.

#### αβ lobes

Manipulating *Dop1R1* expression in the αβ lobes within the Janelia background did not significantly alter social spacing in either sex (two-way ANOVA: effect of genotype, *p* = *0.2479*; effect of genotype × sex interaction, *p = 0.2188*; **Figure 6B**). In females, this conclusion was supported by comparison to the control dataset, as the *Dop1R1-RNAi* construct showed no baseline effect. Accounting for the male-specific baseline effect, however, suggested that males settled farther apart following αβ-lobe *Dop1R1* manipulation, corresponding to a 36% normalized decrease in flies/4BL. Since this effect only emerged after baseline correction, it is tentatively concluded that *Dop1R1* manipulation in the αβ lobes may increase social spacing in males.

#### α’β’ lobes

Manipulating *Dop1R1* expression in the α’β’ lobes within the Janelia background increased social spacing in females but not males (two-way ANOVA: effect of genotype, *p* = *0.0008*; effect of genotype × sex interaction, *p = 0.0041*; **Figure 6C**). Post-hoc Šidak’s tests provided further evidence that female *α’β’>Dop1R1* flies were farther than controls (Šidak’s test: *p = 0.0001*). Comparison to the control dataset reinforced this conclusion, resulting in a normalized 31% decrease in flies/4BL after baseline correction in females. Accounting for baseline effects suggested that males also settle farther apart relative to controls, corresponding to a normalized 56% decrease in flies/4BL. Since this effect only emerged after baseline correction, it is tentatively concluded that *Dop1R1* manipulation in the α’β’ lobes may increase spacing in males.

#### γ lobes

Manipulating *Dop1R1* expression in the γ lobes within the Janelia background did not significantly alter social spacing in either sex (two-way ANOVA: effect of genotype, *p* = *0.6947*; effect of genotype × sex interaction, *p = 0.0462*; **Figure 6D**). Comparison to the control dataset supported this conclusion in females, but accounting for the male-specific baseline effect indicated that males settled farther apart following γ-lobe *Dop1R1* manipulation, corresponding to an 87% normalized decrease in flies/4BL. Since this effect only emerged after baseline correction, it is tentatively concluded that *Dop1R1* manipulation in the γ lobes may increase social spacing in males.

#### Summary

In males, *Dop1R1* manipulation in all three lobes may cause flies to settle farther apart relative to controls, recapitulating the possible increase in social spacing observed at the whole-MB level. In females, manipulating *Dop1R1* expression in the α’β’ lobes increased spacing relative to controls, which was unexpected given that *Dop1R1* showed no influence on social space behavior at the whole-MB level.

### Dop1R2 manipulation decreases male spacing in αβ and α′β′ and decreases female spacing in αβ and γ lobes

#### Baseline effect of *Dop1R2-RNAi* construct

*Dop1R2-RNAi* males did not differ in spacing relative to *attP2* controls. In contrast, females settled farther apart, indicating a female-specific baseline effect of the RNAi construct even in the absence of a Gal4 driver (see Section 4.1.3 for details; **Figure 7A**).

**Figure 7.**
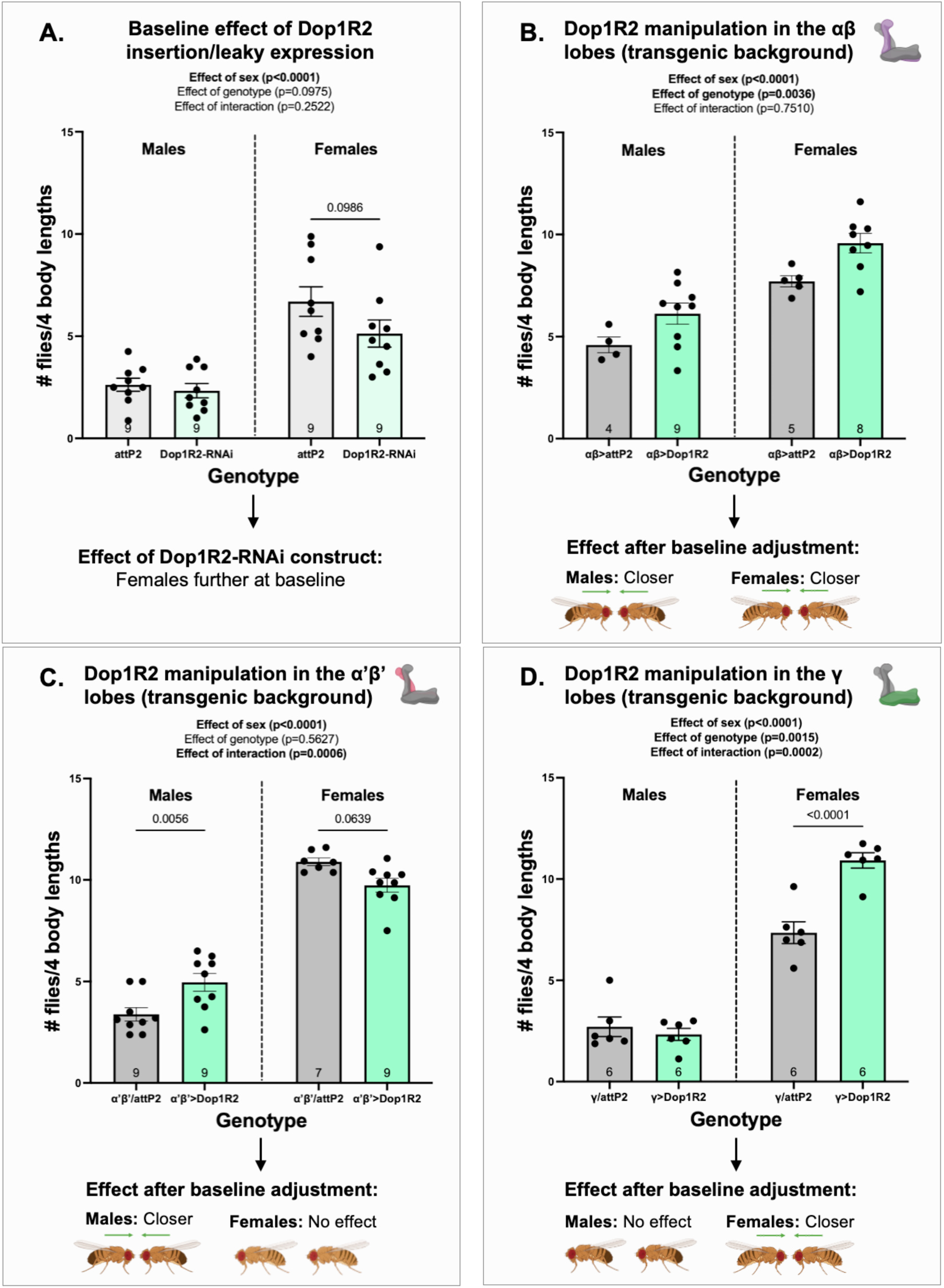
*Dop1R2* manipulation decreases spacing in lobe– and sex-specific ways. Bars show the mean number of flies within four body lengths (± SEM) for each genotype, with lower values indicating farther/increased spacing. Each dot represents one chamber of 12-18 flies (N=4-9 chambers per genotype). Data were analyzed by two-way ANOVA (factors: sex and genotype); graphs display all main and interaction effects, with post-hoc Šidak’s p-values included when relevant for interpretation. Comparison of *Dop1R2-RNAi* flies with *attP2* controls in the control dataset **(A)** reveals a baseline effect of the RNAi construct, with females settling farther apart even in the absence of a Gal4 driver. Manipulating *Dop1R2* expression within the αβ **(B)**, α’β’ **(C)**, and γ **(D)** lobes reveals lobe-specific contributions to social spacing within each sex after baseline correction. *Dop1R2* decreased social spacing in males when manipulated in the αβ and α’β’ lobes, whereas manipulation in the αβ and γ lobes decreased spacing in females.

#### αβ lobes

Manipulating *Dop1R2* expression in the αβ lobes within the Janelia background decreased social spacing in both sexes, with flies settling closer together relative to controls (two-way ANOVA: effect of genotype, *p* = *0.0036*; effect of genotype × sex interaction, *p = 0.7510*; **Figure 7B**). Comparison to the control dataset supported this conclusion in both sexes, with corresponding normalized increases in flies/4BL of 40% in males and 44% in females.

#### α’β’ lobes

Manipulating *Dop1R2* expression in the α’β’ lobes within the Janelia background produced opposite effects across sexes: males showed decreased spacing, whereas females displayed a trend toward increased spacing (two-way ANOVA: effect of genotype, *p* = *0.5627*; effect of genotype × sex interaction, *p = 0.0006*; **Figure 7C**). Since the sex effects were similar in magnitude but opposite in direction, the overall genotype effect was not significant, reflecting the offsetting nature of these responses. Post-hoc Šidak’s tests, however, confirmed that males settled closer together (*p = 0.0056*) and supported the trend toward increased spacing in females (*p = 0.0639*). Comparison to the control dataset reinforced the conclusion that males display decreased spacing following *Dop1R2* manipulation in the α’β’ lobes, with a normalized 55% increase in flies/4BL after baseline correction. In contrast, the increased spacing in females mirrored the baseline pattern in the control dataset, leading to no net effect of α’β’-lobe *Dop1R2* manipulation.

#### γ lobes

Manipulating *Dop1R2* expression in the γ lobes within the Janelia background decreased social spacing in females but not males (two-way ANOVA: effect of genotype, *p* = *0.0015*; effect of genotype × sex interaction, *p = 0.0002*; **Figure 7D**). Post-hoc Šidak’s tests provided further evidence that female γ*>Dop1R2* flies were closer than controls (Šidak’s test: *p < 0.0001*). Comparison to the control dataset reinforced this conclusion, with females showing a normalized 69% increase in flies/4BL after baseline correction. In males, γ-lobe *Dop1R2* manipulation had no effect on social spacing, and this was consistent with the absence of a baseline effect in the control dataset.

#### Summary

In males, *Dop1R2* manipulation in both the αβ and α’β’ lobes caused flies to settle closer together relative to controls, recapitulating the decrease in social spacing observed at the whole-MB level. In females, *Dop1R2* manipulation in the αβ and γ lobes also decreased spacing, consistent with the effect seen when the receptor is manipulated across the entire MB structure.

### Dop2R manipulation decreases male spacing in all lobes and may decrease female spacing in the γ lobe

#### Baseline effect of *Dop2R-RNAi* construct

*Dop2R-RNAi* males did not differ in spacing relative to *attP2* controls. In contrast, females settled farther apart, indicating a female-specific baseline effect of the RNAi construct even in the absence of a Gal4 driver (see Section 4.1.4 for details; **Figure 8A**).

**Figure 8.**
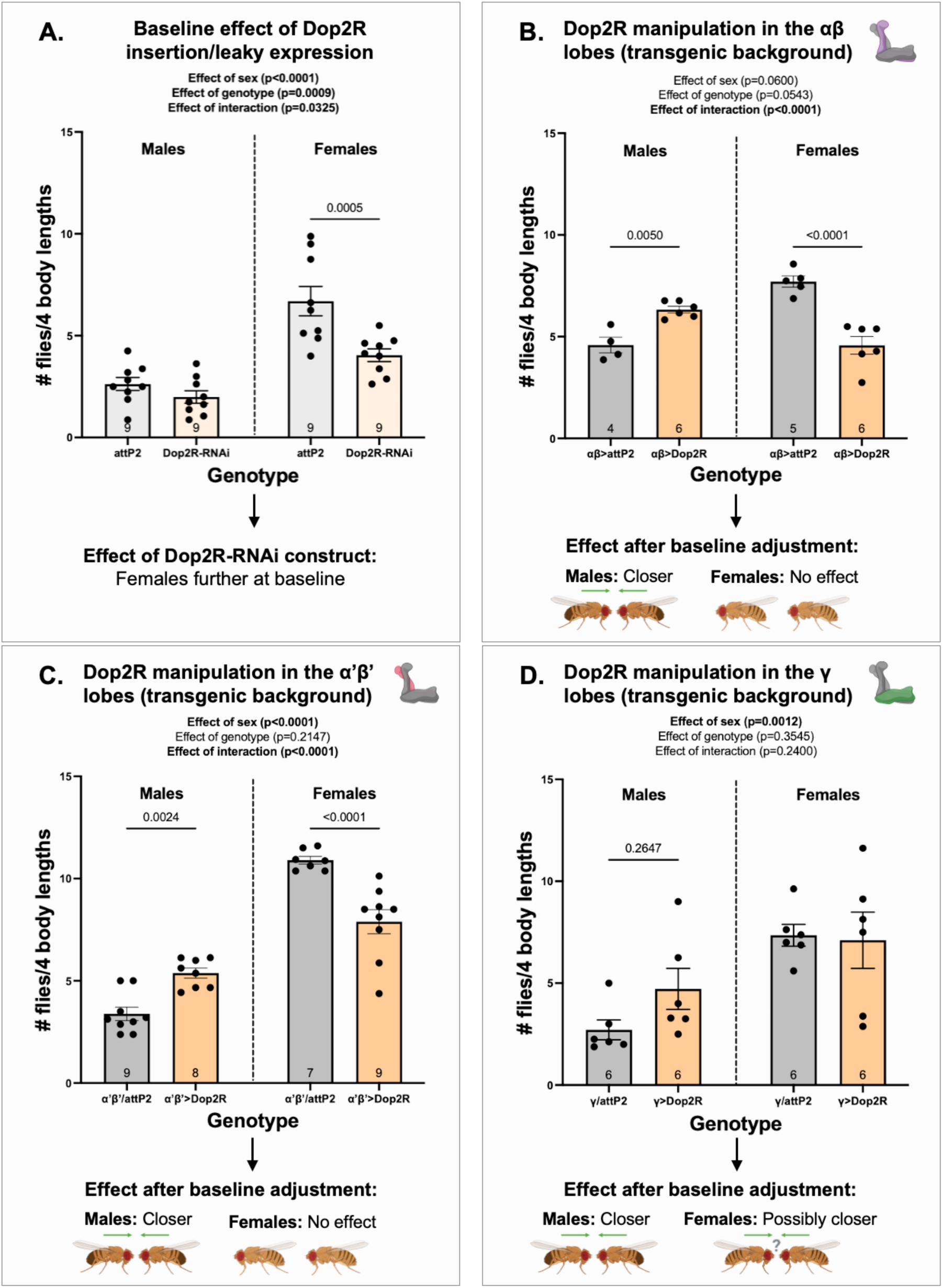
*Dop2R* manipulation decreases spacing in lobe– and sex-specific ways. Bars show the mean number of flies within four body lengths (± SEM) for each genotype, with lower values indicating farther/increased spacing. Each dot represents one chamber of 12-18 flies (N=4-9 chambers per genotype). Data were analyzed by two-way ANOVA (factors: sex and genotype); graphs display all main and interaction effects, with post-hoc Šidak’s p-values included when relevant for interpretation. Comparison of *Dop2R-RNAi* flies with *attP2* controls in the control dataset **(A)** reveals a baseline effect of the RNAi construct, with females settling farther apart even in the absence of a Gal4 driver. Manipulating *Dop2R* expression within the αβ **(B)**, α’β’ **(C)**, and γ **(D)** lobes reveals lobe-specific contributions to social spacing within each sex after baseline correction. *Dop2R* decreases social spacing in males when manipulated in any of the three lobes, whereas manipulation in the γ lobes may decrease spacing in females.

#### αβ lobes

Manipulating *Dop2R* expression in the αβ lobes within the Janelia background produced opposite effects across sexes: males exhibited decreased social spacing, whereas females showed increased spacing (two-way ANOVA: effect of genotype, *p* = *0.0543*; effect of genotype × sex interaction, *p < 0.0001*; **Figure 8B**). Post-hoc Šidak’s tests supported these effects in both males (*p = 0.0050*) and females (*p < 0.0001*). In males, comparison to the control dataset reinforced the conclusion that flies settle closer together following αβ-lobe *Dop2R* manipulation, with a normalized 52% increase in flies/4BL after baseline correction. However, the increased spacing observed in females mirrored the baseline pattern in the control dataset, resulting in no overall effect of αβ-lobe *Dop2R* manipulation after baseline correction.

#### α’β’ lobes

Manipulating *Dop2R* expression in the α’β’ lobes within the Janelia background produced opposite effects across sexes: males exhibited decreased spacing, whereas females showed increased spacing (two-way ANOVA: effect of genotype, *p* = *0.2147*; effect of genotype × sex interaction, *p < 0.0001*; **Figure 8C**). Since the sex effects were similar in magnitude but opposite in direction, the overall genotype effect was not significant, reflecting the offsetting nature of these responses. Post-hoc Šidak’s tests, however, confirmed that males settled closer (*p = 0.0024*) while females were farther apart (*p < 0.0001*). Comparison to the control dataset reinforced the conclusion that α′β′-lobe *Dop2R* manipulation decreases spacing in males, resulting in a normalized 78% increase in flies/4BL after baseline correction. In contrast, the increased spacing in females mirrored the baseline pattern in the control dataset, leading to no net effect of α’β’-lobe *Dop2R* manipulation.

#### γ lobes

Manipulating *Dop2R* expression in the γ lobes within the Janelia background did not significantly alter social spacing in either sex, though males showed a trend toward decreased spacing (two-way ANOVA: effect of genotype, *p* = *0.3545*; effect of genotype × sex interaction, *p = 0.2400*; **Figure 8D**). Post-hoc Šidak’s tests did not strongly support this trend in males (*p = 0.2647*), likely reflecting variability in the data. However, comparison to the control dataset supported the conclusion that males settle closer following γ-lobe *Dop2R* manipulation, resulting in a normalized 66% increase in flies/4BL. Accounting for the female-specific baseline effect indicated that females may also settle closer with this manipulation, with a 32% normalized increase in flies/4BL. Since this effect only emerged after baseline correction, it is tentatively concluded that *Dop2R* manipulation in the γ lobes may decrease social spacing in females.

#### Summary

In males, *Dop2R* manipulation in all three lobes caused flies to settle closer together relative to controls, recapitulating the decrease in social spacing observed at the whole-MB level. In females, manipulating *Dop2R* expression in the γ lobes may decrease spacing relative to controls, consistent with the effect seen when the receptor is manipulated across the entire MB structure.

## Discussion

### Examining patterns in the data

All four dopamine receptors contributed to social spacing in the Janelia background, but their roles were highly distinct both between and within sexes (**Table 1**). Examining receptor function across mushroom body (MB) lobes revealed that no two receptors produced the same pattern of effects when male and female data were considered together, underscoring the specialized contribution of each receptor to this behavior.

**Table 1.** Summary of social spacing effects of *DopR-RNAi* across drivers and sexes. These tables summarize the effects of *driver>DopR-RNAi* manipulations on social spacing after correcting for baseline behavioral effects observed in the control dataset. Each row corresponds to a receptor and each column to a driver. Cells labeled “farther” (red) indicate increases in spacing (fewer flies/4BL, flies farther apart), while cells labelled “closer” (green) indicate decreases in spacing (more flies/4BL, flies closer together). Lighter red or green cells labeled “possibly further” or “possibly closer” indicate effects that emerge only after baseline correction and should be interpreted with caution. White cells indicate no supported effect. Normalized percent changes are provided within each cell. Male and female tables are presented separately.

| Receptor | MB>RNAi | $\alpha\beta$ >RNAi | $\alpha'\beta'$>RNAi | $\gamma$ >RNAi |
| --- | --- | --- | --- | --- |
| DopEcR | Possibly farther<br>↓ 110%* | Possibly farther<br>↓ 65% | No effect<br>↑ 0.3% | Possibly farther<br>↓ 52% |
| Dop1R1 | Possibly farther<br>↓ 78% | Possibly farther<br>↓ 36% | Possibly farther<br>↓ 56% | Possibly farther<br>↓ 87% |
| Dop1R2 | Closer<br>↑ 68% | Closer<br>↑ 40% | Closer<br>↑ 55% | No effect<br>↓ 3% |
| Dop2R | Closer<br>↑ 68% | Closer<br>↑ 52% | Closer<br>↑ 78% | Closer<br>↑ 66% |

**Females**
| Receptor | MB>RNAi | $\alpha\beta$ >RNAi | $\alpha'\beta'$>RNAi | $\gamma$ >RNAi |
| --- | --- | --- | --- | --- |
| DopEcR | No effect<br>↑ 2% | No effect<br>↓ 2% | Farther<br>↓ 35% | No effect<br>↑ 7% |
| Dop1R1 | No effect<br>↓ 3% | No effect<br>↑ 11% | Farther<br>↓ 31% | No effect<br>↑ 9% |
| Dop1R2 | Closer<br>↑ 41% | Closer<br>↑ 44% | No effect<br>↑ 4% | Closer<br>↑ 69% |
| Dop2R | Closer<br>↑ 49% | No effect<br>↓ 6% | No effect<br>↑ 3% | Possibly closer<br>↑ 32% |

\*The normalized percent change of >100% represents a case where the baseline effect was much larger in magnitude than the experimental effect, mathematically amplifying effect size beyond bounded limits. All values are retained for completeness but should be interpreted in relative terms (direction and magnitude) rather than as biologically literal percentages.

Despite this diversity, a consistent organizational pattern emerged: the receptors grouped into two functional pairs that appear to guide the opposing directional components of social spacing that bias flies either toward or away from conspecifics. Manipulating *DopEcR* or *Dop1R1* consistently increased inter-individual distances. This suggests that both receptors normally promote social approach, as disrupting their expression causes flies to space themselves farther apart. In contrast, manipulating *Dop1R2* or *Dop2R* reliably decreased inter-individual distances. This pattern indicates that these receptors typically support social retreat, with disrupted expression resulting in flies settling closer together than expected.

All four receptors influenced social spacing in at least one MB lobe, with several showing effects across multiple lobes (**Table 1**). In most cases, the lobe-specific effects of each receptor mirror the effect observed when expression is manipulated across the whole MB. The two exceptions are *DopEcR* and *Dop1R1* in females: manipulating either receptor in the α′β′ lobe increases spacing, but similar manipulations in the αβ or γ lobes or across the entire MB show no detectable effect. Although this pattern may appear counterintuitive, it is consistent with a framework in which these receptors exert opposing influences of different magnitudes across the three lobes (**Supp. Figure 9**). It is possible that DopEcR and Dop1R1 produce relatively weak retreat-promoting signals in the αβ and γ lobes and a stronger approach-promoting signal in the α′β′ lobe. Under normal conditions, these opposing signals may balance each other. Disrupting expression in a single αβ or γ lobe may not produce a detectable effect due to compensation from the other lobe, whereas combined manipulation could reveal reduced spacing. In contrast, disrupting the α′β′ lobe removes the dominant approach signal, allowing retreat signals to prevail and increasing spacing.

Support for this interpretation comes from a study by Aranda et al. (2017), which showed that reducing *DopEcR* in the αβ and γ lobes together impaired ethanol-induced behavioral sensitization, whereas single-lobe manipulations had no effect. This pattern is consistent with compensatory lobe-specific contributions masking phenotypes when lobes are examined in isolation.

Collectively, all four dopamine receptors broadly contribute to social spacing, with sexually dimorphic effects spanning all three MB lobes. The extensive receptor-, sex-, and lobe-specific influences observed likely reflect the complexity of social spacing and its underlying circuitry, which require the integration of diverse contextual cues to enable continuous positional adjustments in a dynamic social environment.

### Incorporating findings with current literature: towards a circuit model

The widespread involvement of all four dopamine receptors across MB lobes suggests that social spacing reflects distributed processing throughout the MB, rather than signaling localized to specific receptors or regions. This is consistent with the known organisation of the MB, in which different compartment-specific dopaminergic neurons (DANs) deliver distinct contextual signals to kenyon cells (KCs) in compartments across the αβ, α′β′, and γ lobes (Aso et al., 2014a). These DAN signals modulate KC output at *en passant* synapses along KC axons, altering how sensory inputs received at the calyx are relayed to downstream MB circuits (**Supp. Figure 3**).

Social spacing is a highly context-dependent behavior influenced by multiple factors, including time of day, mating status, and prior social experience, among others (McNeil et al., 2012; Simon et al., 2012; Yost et al., 2020). Since these cues are diverse, they are likely encoded by multiple DAN types that collectively span the MB. In this way, distributed dopaminergic input provides a basis for integrating varied contextual information across MB subregions.

We propose that this distributed dopaminergic input is then differentially weighted at the level of KCs through region-specific variation in dopamine receptor expression. Each receptor type differs in its response to dopamine and exhibits distinct expression patterns across MB lobes and compartments (Han et al., 1996; Kim et al., 2003; Croset et al., 2018; Karam et al., 2020; Kondo et al., 2020). This heterogeneity means that the same dopaminergic input can be differentially weighted depending on the local receptor complement at a given KC membrane. For example, a KC membrane enriched for excitatory receptors such as Dop1R1 would be expected to exhibit stronger depolarising responses to dopamine than a membrane with higher proportions of inhibitory Dop2R, resulting in graded modulation of KC output that can differ across compartments. KCs are cholinergic, and thus these differences in excitability translate into differences in acetylcholine release (Barnstedt et al., 2016).

Graded KC output is then transmitted to mushroom body output neurons (MBONs), a population of 34 neurons that each receive input from KCs within specific, compartment-restricted regions of the MB (Aso et al., 2014a). Evidence from other MB-dependent behaviors suggests that MBONs operate within an opponent-process framework, in which distinct populations drive opposing behavioral biases. This principle is well established in olfactory learning and sleep regulation, where MBON groups promote opposite behaviors depending on their transmitter identity (Aso et al., 2014b; Sitaraman et al., 2015).

Extending this framework to social spacing, we propose that MBONs encode either social approach or social retreat biases, with individual neurons additionally capable of a neutral state when KC drive is insufficient to reach activation threshold (**Figure 9**). In this model, MBON identity determines the valence of the signal and the magnitude of KC input determines signal strength. Strong KC input generates robust approach– or retreat-biased responses, and weaker input produces proportionally attenuated or neutral MBON activity (**Figure 10A**).

**Figure 9.**
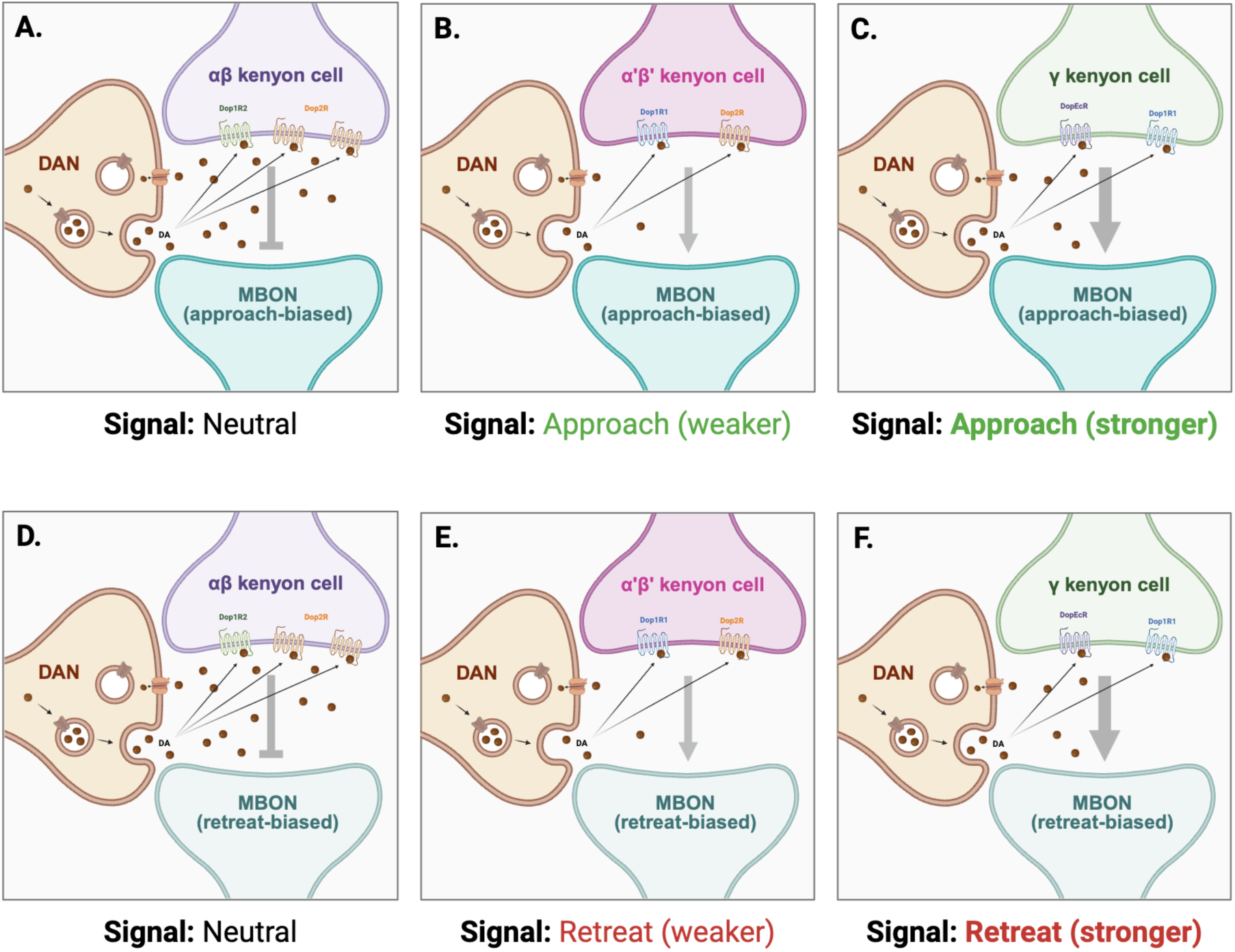
Spacing-relevant signals encoded by mushroom body output neurons. Dopamine receptors on Kenyon cell (KC) membranes respond to modulatory input from dopaminergic neurons (DANs), which is integrated with sensory information entering at the calyx. The resulting KC output depends on both the incoming signals and the specific complement of receptors expressed by the KC. KC axons synapse onto mushroom body output neurons (MBONs), and the identity of the postsynaptic MBON determines the type of signal passed down the social spacing pathway. In the proposed model, MBONs can be either approach-biased **(A-C)** or retreat-biased **(D-F)**. For approach-biased MBONs, weak KC output below the MBON’s activation threshold produces a neutral signal **(A)**, whereas progressively stronger KC outputs generate increasingly strong approach-biased signals **(B, C)**. For retreat-biased MBONs, weak KC output similarly produces a neutral signal **(D)**, while stronger KC outputs yield progressively stronger retreat-biased signals **(E, F)**. Neuron and receptor combinations are illustrative and not intended to represent empirically validated pairings. Image created in BioRender.

**Figure 10.**
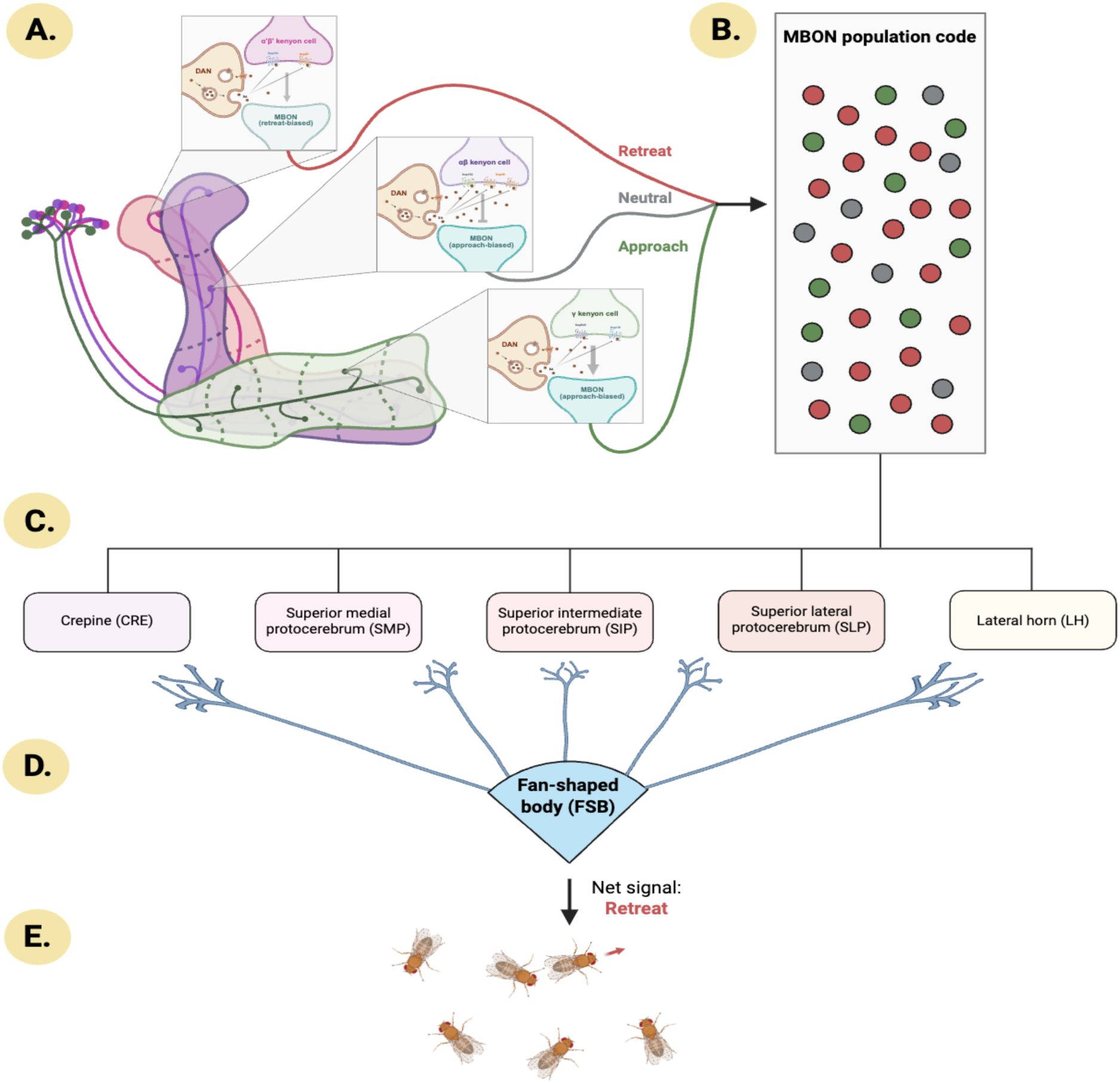
Proposed integrative model of social space circuitry in Drosophila. **A.** Kenyon cell (KC) output varies across mushroom body (MB) lobes and compartments due to differences in dopaminergic input and dopamine receptor expression. KCs synapse onto mushroom body output neurons (MBONs), which generate neutral, approach-biased, or retreat-biased signals of varying strength. **B.** The combined output of all 34 MBONs forms an MBON population code that determines the net approach-retreat bias underlying spacing decisions. **C.** MBONs send axonal projections to five neuropils in the Drosophila brain. **D.** Within these neuropils, MBONs converge on neurons that project to the fan-shaped body (FSB), a sensorimotor hub of the central complex. **E.** The FSB transforms the MBON population code into the positional adjustments that establish and maintain preferred social spacing. Image created in BioRender.

We propose that the individual MBON signals described above can be integrated into a population code in which the collective activity of all 34 MBONs determines a net bias toward social approach, social retreat, or no change in position (**Figure 10B**). This framework is supported by evidence from other MB-dependent behaviors showing that MBON effects can be additive, along with the neuroanatomical finding that MBONs converge on many shared postsynaptic targets (Aso et al., 2014a; Aso et al., 2014b). Applying an MBON population code to social spacing explains how small changes in contextual input or receptor-mediated modulation can be translated into appropriately scaled positional adjustments.

MBON outputs are known to be transmitted to five downstream neuropil regions: the crepine (CRE); the superior medial protocerebrum (SMP); the superior intermediate protocerebrum (SIP); the superior lateral protocerebrum (SLP); and the lateral horn (LH) (**Figure 10C**) (Aso et al., 2014a). Within these neuropils, the axons of many MBONs converge onto the dendrites of neurons that in turn project to the fan-shaped body (FSB) (**Figure 10D**).

As a central complex structure with well-established roles in sensorimotor integration, the FSB is well positioned to integrate these convergent MBON-derived signals (Aso et al., 2014a). We propose that the FSB acts as the downstream readout of the MBON population code, transforming the net approach-retreat bias into motor commands that regulate movement toward or away from neighboring individuals (**Figure 10E**). In this way, the connectivity between MBONs and the FSB provides a plausible pathway through which distributed MB computations are translated into the fine-scale positional adjustments that underlie social spacing.

Together, this framework links compartment-specific DAN input, receptor-dependent KC responses, and MBON population coding into a unified circuit-level model that accounts for the distributed and context-dependent nature of receptor contributions we observed in social spacing.

### Explaining sexual dimorphism with the model

In addition to explaining widespread receptor– and lobe-specific contributions to social spacing, our proposed circuit model provides a mechanistic framework for the sexually dimorphic effects we observed. Within this framework, sex-specific variation in KC-level connectivity and dopamine receptor expression profiles may alter patterns of receptor involvement without requiring changes to the underlying circuit architecture. We first consider variation in KC-level connectivity, followed by differences in dopamine receptor expression.

Although DAN and MBON projection patterns to individual compartments are fixed, the KC-level wiring within those compartments may not be (Aso et al., 2014a). Variability in which KCs receive input from particular DANs, or which KCs provide output to specific MBONs, could reshape the synaptic landscape by altering the number and distribution of DAN-KC-MBON synapses. In compartments with multiple DANs or MBONs, such differences additionally determine which combinations of synaptic partners a given KC engages with, expanding the possible configurations of local synapses and altering which dopaminergic cues individual KCs can access. These shifts in local microcircuit topology could, in turn, modify which receptors appear behaviorally relevant when expression is experimentally manipulated.

One potential source of variation in KC connectivity is the sexually dimorphic *fruitless* (*fru*) transcript, which encodes a sex-specific transcription factor and has been linked to dimorphic neuronal wiring (Cachero et al., 2010). Although the functional Fru protein is only expressed in males, *fru* transcripts are expressed in the MB of both sexes (**Supp. Figure 10, 11**) (Palmateer et al., 2023). At the whole-brain level, *fru*+ neurons exhibit widespread sex-specific connectivity patterns, with males often showing denser arborization (Cachero et al., 2010; Palmateer et al., 2023). While most work has focused on global *fru*+ populations, the MB displays anatomical differences between sexes that are indicative of dimorphic connectivity, including increased fibre number in females and enlarged medial γ lobe tips in males (Technau, 1984; Cachero et al., 2010). Together, these findings support the idea that KC-level connectivity may differ between sexes, providing a plausible mechanism by which shared circuit architecture could give rise to sex-specific patterns of dopamine receptor involvement in social spacing.

A second parameter that could alter receptor involvement is dopamine receptor expression profiles. Changes in the location or the density of receptor expression across MB lobes would influence where each receptor is positioned to detect dopaminergic signals and how it interacts with other receptors that may be at the same KC. Even when spatial localization is unchanged, variation in receptor density can meaningfully alter the balance of excitatory and inhibitory drive. For instance, high Dop2R expression at a synapse that also contains Dop1R1 may counteract or even override Dop1R1-mediated excitation. In contrast, lower Dop2R levels at the same synapse may not provide sufficient inhibition for Dop2R loss to have a detectable behavioral effect. In this way, receptor expression patterns critically shape which receptors contribute to social spacing and where within the MB those contributions arise.

The distinct expression profiles of the four dopamine receptors have not been characterized in a sex-specific manner within the MB. Nonetheless, reported differences in average receptor expression between male and female heads suggest that expression levels may vary across sexes at the whole-brain level (Leader et al., 2018). In addition, many genes showing sex-biased expression are associated with *fru*+ neuron populations, which are present in the MB (Palmateer et al., 2023; Dähn & Wagner, 2025). Together, these observations support the possibility that dopamine receptor expression within the MB differs between males and females, providing an additional mechanism through which receptor contributions to social spacing may diverge despite a shared core circuit.

### Genetic background effects

A separate study by Robinson et al. (2024) used the same MB driver and RNAi effector lines to examine dopamine receptor contributions to social spacing, with one key difference: the MB driver was outcrossed into a Canton-Special (CS) background rather than maintained in the original Janelia background. This study reported MB-specific receptor effects that differed from some of those observed here, indicating that the underlying circuitry is sensitive to genetic context. This interpretation is consistent with prior findings that social spacing itself varies across genetic backgrounds, with some Drosophila strains forming tighter groups and others settling farther apart (McNeil et al., 2012; Jiménez-Padilla et al., 2024).

The same two circuit parameters proposed to account for sex differences in receptor contributions may also explain background-specific effects. Genetic background has been shown to influence KC morphology in the MB, with evidence that these anatomical differences reflect variation in underlying connectivity. For example, Zwarts et al. (2015) reported substantial strain-dependent differences in the length and width of αβ lobe KCs and suggested that these changes likely reflect alterations in the local synaptic landscape. Genetic background is also known to shape transcriptional profiles in *Drosophila*, with commonly used wild-derived strains exhibiting marked differences in gene expression (Dworkin et al., 2009; Zarubin et al., 2020).

Although dopamine receptor expression in the MB has not been directly compared between Janelia and CS backgrounds, differences in expression are plausible. Together, variation in KC-level connectivity and receptor expression across genetic backgrounds may therefore contribute to the differences in receptor involvement observed across studies.

### Future directions

We propose an approach for analysing genetic data that accounts for baseline effects of RNAi constructs without requiring lengthy outcrossing schemes. Nonetheless, outcrossing driver lines into a defined genetic background to generate matched non-driver controls would allow each *DopR-RNAi* line to be assessed in a fully controlled genetic context. In addition, this study did not distinguish between developmental and adult-stage effects of receptor manipulation; the use of a temperature-sensitive Gal80 system (Gal80^TS^) to restrict RNAi expression to adulthood would therefore help isolate receptor function from developmental compensation or long-term circuit adaptation. Future studies should also consider the relative contributions of 20-hydroxyecdysone and dopamine to DopEcR-mediated effects in social spacing, given that both ligands can bind this dual-ligand receptor (Karam et al., 2020).

The proposed social space model generates several testable predictions about the circuit mechanisms underlying social spacing, and future work should explore these avenues. Selective manipulation of individual DANs would allow compartment-level probing of contextual inputs, while targeted perturbation of specific MBONs could clarify the relative contributions of MB compartments and directly test the proposed opponent organization and population coding framework. Disrupting neurons of the fan-shaped body would help define its role as a downstream structure in the social spacing circuit. Additional insight into receptor– and lobe-specific contributions could be obtained through a double manipulation approach, in which two receptors or two MB lobes are simultaneously targeted.

The context dependence of the proposed circuitry can also be tested by altering the intrinsic sexual identity of MB neurons to determine whether sex-specific effects are reversed, and by outcrossing driver lines into additional genetic backgrounds followed by replication of receptor manipulations in these strains. In parallel, KC-level connectivity and dopamine receptor expression profiles could be quantified across sexes and backgrounds to identify differences that may underlie the distinct behavioral phenotypes observed. Collectively, these experimental approaches provide a clear roadmap for testing and extending the proposed social space model.

### Significance

By manipulating dopamine receptor expression within the MB, this work establishes a previously uncharacterized functional link between dopaminergic signaling and the MB in the regulation of social spacing. Dopaminergic contributions to social spacing are highly context dependent, varying across receptor type, lobe, sex, and genetic background. To account for these findings, we propose an integrative circuit model in which MB-localized dopaminergic signals are flexibly weighted through region-specific receptor expression and transformed via downstream MBON pathways to generate spacing decisions. This framework incorporates adjustable circuit parameters that can shift across biological and genetic contexts, providing a mechanistic explanation for the diverse receptor– and lobe-specific effects observed and generating a range of testable predictions across multiple circuit levels.

Beyond Drosophila, these findings may have broader relevance. Dopaminergic signaling is strongly conserved across phyla, and fundamental social behaviors such as social spacing are likely shaped by evolutionary ancient circuit motifs (Toth & Robinson, 2007; Sokolowski et al., 2010; Reaume & Sokolowski, 2011; Yamamoto & Vernier, 2011; Goodson & Kingsbury, 2013; Li et al., 2020). As such, this work may provide insight into general principles governing how neuromodulatory systems regulate social behavior in other organisms. More broadly, the context-dependent effects uncovered here highlight both the importance of experimental specificity and the inherent complexity of biological systems.

## Acknowledgments

This study would not have been possible without the continued funding received by Western’s Biotron Facility and the support offered there by the staff: we are very thankful to Karen Nygard for her technical and molecular help in immunofluorescence protocol establishment and microscopy within the integrated microscopy unit. We also want to thank the Insect Suite within the Biotron Facility of Western University, for housing our flies in climate-controlled walk-in rooms.

We are immensely grateful to the essential Flybase (OZTURK-COLAK *et al*. 2024)– NHGRI Awards U41HG000739 and U24HG010859), and to the Bloomington Drosophila Stock Center (NIH P40OD018537) for the invaluable resources they provide to the scientific community.

## Ethics Statement

University of Western Ontario’s Animal Care Committee and Ontario Provincial and Federal regulatory bodies specify that ethical considerations do not apply to the study of invertebrate animals including insects such as *Drosophila melanogaster*.

## Data availability statement

The datasets analysed for this study will be deposited in the Dryad repository

## Authors contribution

Based on Oxford Academic (Genetics journal)

MRE: Conceptualization, Supervision, Project Administration, Investigation, Formal Analysis, Methodology, Resources, Funding Acquisition, Writing – Original Draft Preparation, and Writing – Review & Editing.

AFS: Conceptualization, Supervision, Project Administration, Formal Analysis, Resources, Funding Acquisition, and Writing – Review & Editing.

## Supplemental Table and Figure Captions

**Supplemental Table 1.** Key properties of the four dopamine receptors in Drosophila. For each receptor, the chromosomal location, receptor class, intracellular signaling cascades, physiological effect on neuronal excitability, subcellular localization, and unique behavioral roles are summarized. Chromosomal locations refer to the X chromosome or the left (3L) and right (3R) arms of chromosome 3 in the Drosophila genome. Subcellular localization includes synaptic position (presynaptic vs. postsynaptic) as well as localization within the mushroom bodies. Only behavioral roles that are unique to each receptor are listed. Chromosomal locations were obtained from FlyBase (https://flybase.org/). *DopEcR also binds the ecdysteroid 20-hydroxyecdysone (20E), activating MAPK/ERK signalling.

| Receptor | Chromosomal Location | Class | Signalling Cascade | Physiological Effect | Subcellular Localization (MB) | Unique Behavioural Roles | References |
| --- | --- | --- | --- | --- | --- | --- | --- |
| **DopEcR\*** | 3L | non-canonical | cAMP/PKA (↑ cAMP)<br>P13K/Akt<br>MAPK/ERK* | excitatory | postsynaptic (KC axon initiation segment) | sex pheromone response, appetite, adaptation to stress, courtship memory, ethanol-induced behavioural sensitization | Inagaki et al., 2012; Abrieux et al., 2014; Aranda et al., 2017; Karam et al., 2020; Petruccelli et al., 2020; Deng et al., 2019 |
| **Dop1R1** | 3R | D1 | cAMP/PKA (↑ cAMP) | excitatory | postsynaptic (uniform across KC membrane) | long-term memory, male courtship learning, locomotion | Keleman et al., 2012; Karam et al., 2020; Silva et al., 2020 |
| **Dop1R2** | 3R | D1 | cAMP/PKA (↑ cAMP) | excitatory | postsynaptic (anterior peduncle and distal KC axon) | forgetting, male courtship drive, female song response | Pimentel et al., 2016; Zhang et al., 2016; Karam et al., 2020; Yamakoshi et al., 2025 |
| **Dop2R** | X | D2 | cAMP/PKA (↓ cAMP) | inhibitory | presynaptic (DANs), postsynaptic (uniform across KC membrane) | food-seeking behaviour, olfactory associative learning, aversive learning, anesthesia-resistant memory | Scholz-Kornehl & Schwärzel, 2016; Karam et al., 2020 |

**Supplemental Table 2.**
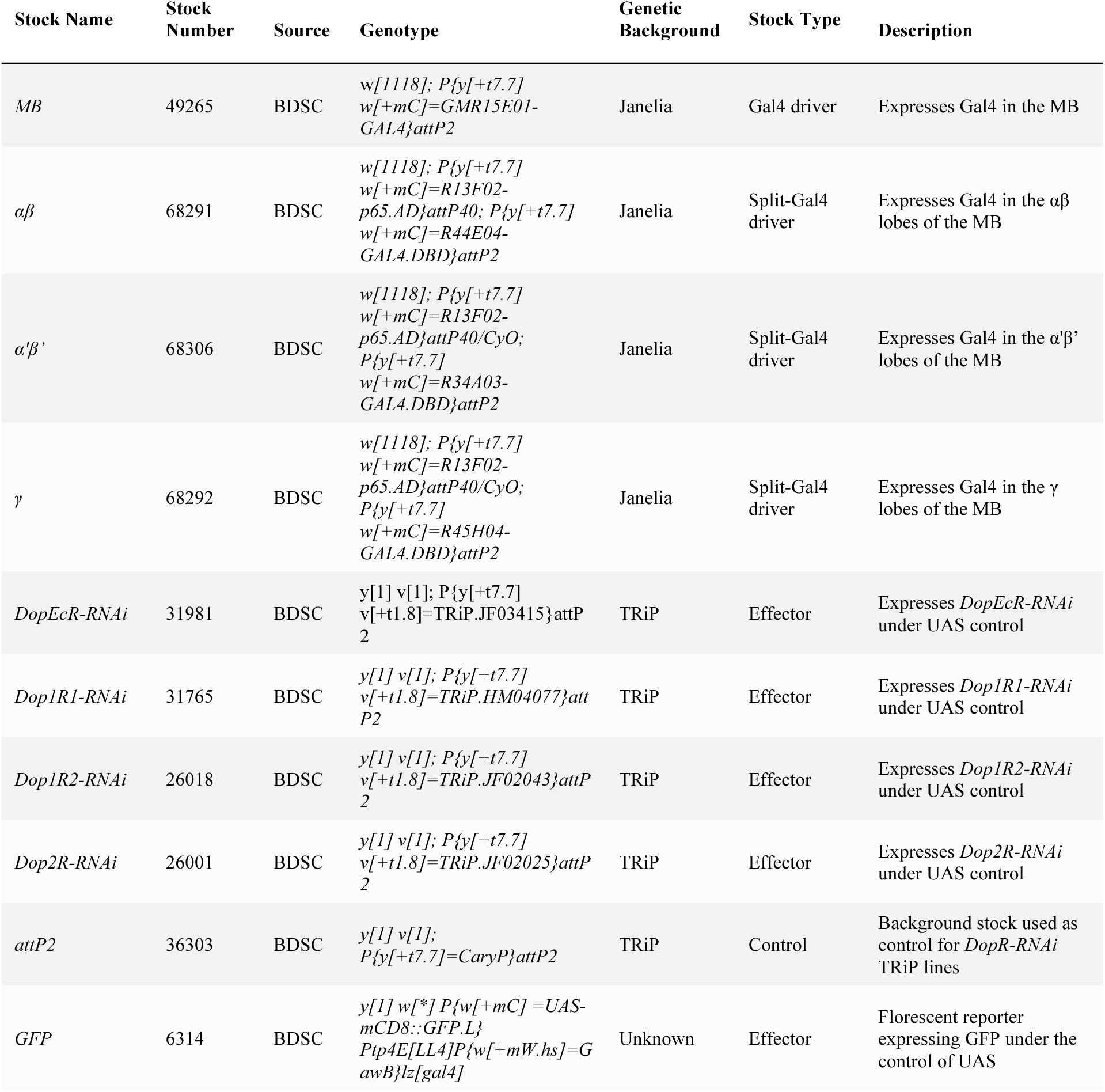
Parental Drosophila stocks used to generate experimental fly lines. Stock names were created for simplicity in text. All stocks are obtained from Bloomington Drosophila Stock Center (BDSC).

| Stock Name | Stock Number | Source | Genotype | Genetic Background | Stock Type | Description |
| --- | --- | --- | --- | --- | --- | --- |
| *MB* | 49265 | BDSC | *w[1118]; P{y[+t7.7]*<br>*w[+mC]=GMR15E01-*<br>*GAL4}attP2* | Janelia | Gal4 driver | Expresses Gal4 in the MB |
| $\alpha\beta$ | 68291 | BDSC | *w[1118]; P{y[+t7.7]*<br>*w[+mC]=R13F02-*<br>*p65.AD}attP40; P{y[+t7.7]*<br>*w[+mC]=R44E04-*<br>*GAL4.DBD}attP2* | Janelia | Split-Gal4 driver | Expresses Gal4 in the $\alpha\beta$ lobes of the MB |
| $\alpha'\beta'$ | 68306 | BDSC | *w[1118]; P{y[+t7.7]*<br>*w[+mC]=R13F02-*<br>*p65.AD}attP40/CyO;*<br>*P{y[+t7.7]*<br>*w[+mC]=R34A03-*<br>*GAL4.DBD}attP2* | Janelia | Split-Gal4 driver | Expresses Gal4 in the $\alpha'\beta'$ lobes of the MB |
| $\gamma$ | 68292 | BDSC | *w[1118]; P{y[+t7.7]*<br>*w[+mC]=R13F02-*<br>*p65.AD}attP40/CyO;*<br>*P{y[+t7.7]*<br>*w[+mC]=R45H04-*<br>*GAL4.DBD}attP2* | Janelia | Split-Gal4 driver | Expresses Gal4 in the $\gamma$ lobes of the MB |
| *DopEcR-RNAi* | 31981 | BDSC | *y[1] v[1]; P{y[+t7.7]*<br>*v[+t1.8]=TRiP.JF03415}attP*<br>*2* | TRiP | Effector | Expresses *DopEcR-RNAi* under UAS control |
| *Dop1R1-RNAi* | 31765 | BDSC | *y[1] v[1]; P{y[+t7.7]*<br>*v[+t1.8]=TRiP.HM04077}att*<br>*P2* | TRiP | Effector | Expresses *Dop1R1-RNAi* under UAS control |
| *Dop1R2-RNAi* | 26018 | BDSC | *y[1] v[1]; P{y[+t7.7]*<br>*v[+t1.8]=TRiP.JF02043}attP*<br>*2* | TRiP | Effector | Expresses *Dop1R2-RNAi* under UAS control |
| *Dop2R-RNAi* | 26001 | BDSC | *y[1] v[1]; P{y[+t7.7]*<br>*v[+t1.8]=TRiP.JF02025}attP*<br>*2* | TRiP | Effector | Expresses *Dop2R-RNAi* under UAS control |
| *attP2* | 36303 | BDSC | *y[1] v[1];*<br>*P{y[+t7.7]=CaryP}attP2* | TRiP | Control | Background stock used as control for *DopR-RNAi* TRiP lines |
| *GFP* | 6314 | BDSC | *y[1] w[\*] P{w[+mC] =UAS-*<br>*mCD8::GFP.L}*<br>*Ptp4E[LL4]P{w[+mW.hs]=G*<br>*awB}lz[gal4]* | Unknown | Effector | Florescent reporter expressing GFP under the control of UAS |

**Supplemental Table 3.** Crosses performed to generate experimental and control lines. Each row represents a genetic crossing scheme, listing the parental driver and effector stocks, the resulting progeny (short name), their genetic background, and the manipulation achieved.

| Driver Parent | Effector Parent | Progeny ID | Genetic Background of Progeny | Manipulation |
| --- | --- | --- | --- | --- |
| *MB* | *attP2* | *MB/attP2* | 50% Janelia, 50% TRiP | Control cross for *MB* driver (no manipulation) |
| *MB* | *DopEcR-RNAi* | *MB>DopEcR* | 50% Janelia, 50% TRiP | *DopEcR* expression manipulated in MB |
| *MB* | *Dop1R1-RNAi* | *MB>Dop1R1* | 50% Janelia, 50% TRiP | *Dop1R1* expression manipulated in MB |
| *MB* | *Dop1R2-RNAi* | *MB>Dop1R2* | 50% Janelia, 50% TRiP | *Dop1R2* expression manipulated in MB |
| *MB* | *Dop2R-RNAi* | *MB>Dop2R* | 50% Janelia, 50% TRiP | *Dop2R* expression manipulated in MB |
| *MB* | *GFP* | *MB>GFP* | 50% Janelia, 50% unknown | GFP expressed in MB to validate driver |
| $\alpha\beta$ | *attP2* | $\alpha\beta/attP2$ | 50% Janelia, 50% TRiP | Control cross for $\alpha\beta$ driver (no manipulation) |
| $\alpha\beta$ | *DopEcR-RNAi* | $\alpha\beta>DopEcR$ | 50% Janelia, 50% TRiP | *DopEcR* expression manipulated in the $\alpha\beta$ lobes of the MB |
| $\alpha\beta$ | *Dop1R1-RNAi* | $\alpha\beta>Dop1R1$ | 50% Janelia, 50% TRiP | *Dop1R1* expression manipulated in the $\alpha\beta$ lobes of the MB |
| $\alpha\beta$ | *Dop1R2-RNAi* | $\alpha\beta>Dop1R2$ | 50% Janelia, 50% TRiP | *Dop1R2* expression manipulated in the $\alpha\beta$ lobes of the MB |
| $\alpha\beta$ | *Dop2R-RNAi* | $\alpha\beta>Dop2R$ | 50% Janelia, 50% TRiP | *Dop2R* expression manipulated in the $\alpha\beta$ lobes of the MB |
| $\alpha\beta$ | *GFP* | $\alpha\beta>GFP$ | 50% Janelia, 50% unknown | GFP expressed in the $\alpha\beta$ lobes of the MB to validate driver |
| $\alpha'\beta'$ | *attP2* | $\alpha'\beta'/attP2$ | 50% Janelia, 50% TRiP | Control cross for $\alpha'\beta'$ driver (no manipulation) |
| $\alpha'\beta'$ | *DopEcR-RNAi* | $\alpha'\beta'>DopEcR$ | 50% Janelia, 50% TRiP | *DopEcR* expression manipulated in the $\alpha'\beta'$ lobes of the MB |
| $\alpha'\beta'$ | *Dop1R1-RNAi* | $\alpha'\beta'>Dop1R1$ | 50% Janelia, 50% TRiP | *Dop1R1* expression manipulated in the $\alpha'\beta'$ lobes of the MB |
| $\alpha'\beta'$ | *Dop1R2-RNAi* | $\alpha'\beta'>Dop1R2$ | 50% Janelia, 50% TRiP | *Dop1R2* expression manipulated in the $\alpha'\beta'$ lobes of the MB |
| $\alpha'\beta'$ | *Dop2R-RNAi* | $\alpha'\beta'>Dop2R$ | 50% Janelia, 50% TRiP | *Dop2R* expression manipulated in the $\alpha'\beta'$ lobes of the MB |
| $\alpha'\beta'$ | *GFP* | $\alpha'\beta'>GFP$ | 50% Janelia, 50% unknown | GFP expressed in the $\alpha'\beta'$ lobes of the MB to validate driver |
| $\gamma$ | *attP2* | $\gamma/attP2$ | 50% Janelia, 50% TRiP | Control cross for $\gamma$ driver (no manipulation) |
| $\gamma$ | *DopEcR-RNAi* | $\gamma>DopEcR$ | 50% Janelia, 50% TRiP | *DopEcR* expression manipulated in the $\gamma$ lobes of the MB |
| $\gamma$ | *Dop1R1-RNAi* | $\gamma>Dop1R1$ | 50% Janelia, 50% TRiP | *Dop1R1* expression manipulated in the $\gamma$ lobes of the MB |
| $\gamma$ | *Dop1R2-RNAi* | $\gamma>Dop1R2$ | 50% Janelia, 50% TRiP | *Dop1R2* expression manipulated in the $\gamma$ lobes of the MB |
| $\gamma$ | *Dop2R-RNAi* | $\gamma>Dop2R$ | 50% Janelia, 50% TRiP | *Dop2R* expression manipulated in the $\gamma$ lobes of the MB |
| $\gamma$ | *GFP* | $\gamma>GFP$ | 50% Janelia, 50% unknown | GFP expressed in the $\gamma$ lobes of the MB to validate driver |

**Supplemental Table 4.**
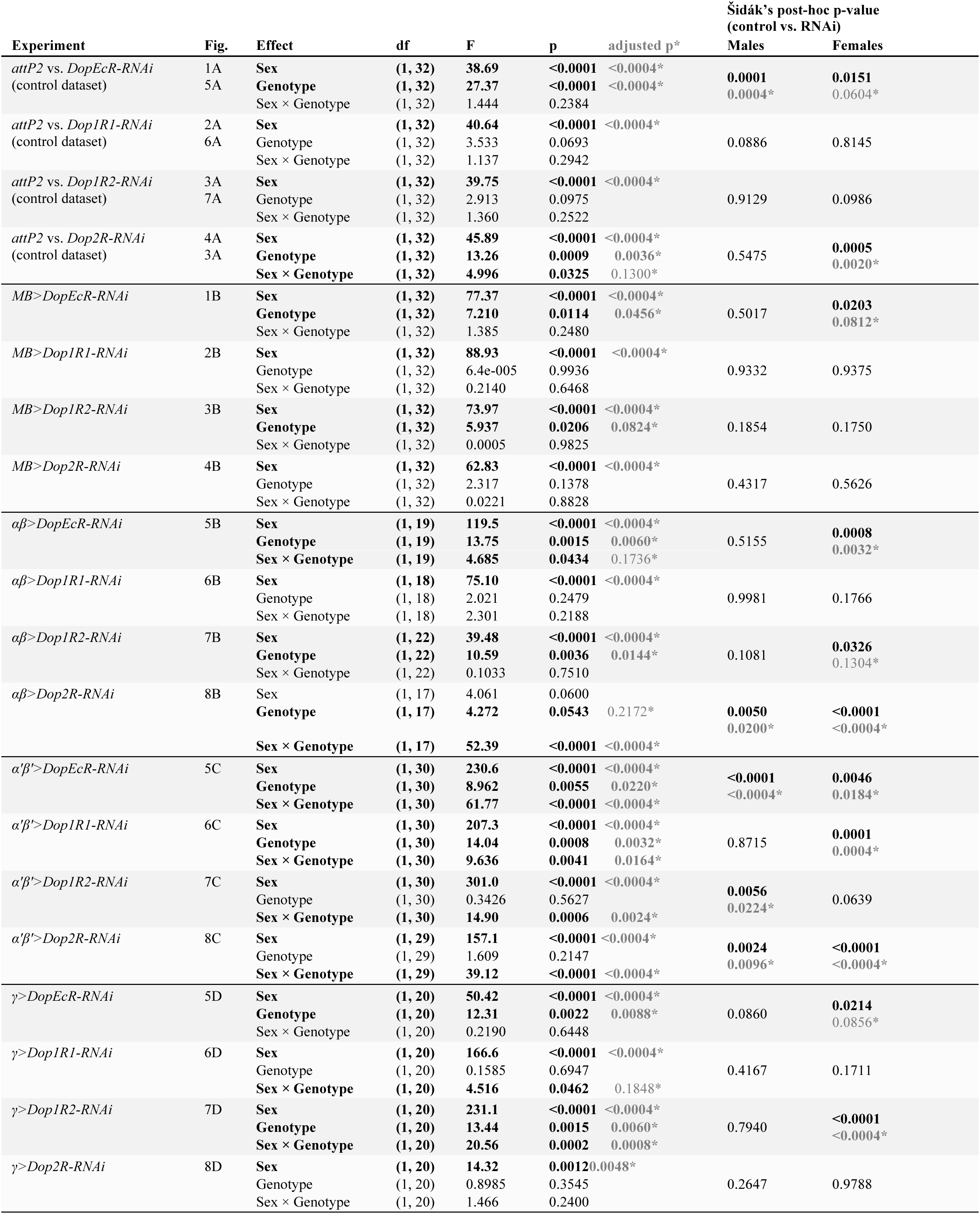
Social space assay statistical outputs. Results of all two-way ANOVA and Šidák post-hoc tests are shown for the control dataset and the four experimental datasets (*MB*, *αβ, α’β’, γ*). All values associated with *p < 0.05* are bolded. *When the same control group was used for multiple comparisons and p < 0.05, Bonferroni-adjusted p-values are reported.

**Supplemental Table 5.** Social space effect sizes and normalized percent change values. Effect sizes were calculated separately for males and females as the difference in mean flies within four body lengths (flies/4BL) between genotypes. Experimental effect sizes were defined as *(driver>RNAi – driver/attP2)*, and baseline effect sizes as *(RNAi – attP2)*. Additive effect sizes were obtained by subtracting the baseline effect size from the corresponding experimental effect size to isolate driver-dependent changes in social spacing. These values were normalized to the mean flies/4BL of the experimental control genotype *(driver/attP2)* to yield a percent change. Bolded values indicate effects in the experimental dataset that remain after control comparison, with red denoting decreased flies/4BL (farther apart) and green denoting increased flies/4BL (closer together). Non-bolded coloured values indicate possible effects emerging only after baseline correction, while black text indicates no evidence of an effect. *The normalized percent change of >100% represents a case where the baseline effect was much larger in magnitude than the experimental effect, mathematically amplifying effect size beyond bounded limits. All values should be interpreted in relative terms rather literal percentages.

| Driver | Sex | Receptor | Experimental Effect Size (Flies/4BL) | Baseline Effect Size (Flies/4BL) | Additive Effect Size (Flies/4BL) | Experimental Control Mean (driver/attP2; Flies/4BL) | Normalized Change (%) |
| --- | --- | --- | --- | --- | --- | --- | --- |
| *MB* | M | DopEcR | +0.92 | +3.53 | -2.60 | 2.37 | ↓ 110%* |
|  |  | Dop1R1 | -0.24 | +1.61 | -1.85 | 2.37 | ↓ 78% |
|  |  | Dop1R2 | +1.31 | -0.30 | +1.61 | 2.37 | ↑ 68% |
|  |  | Dop2R | +0.96 | -0.64 | +1.60 | 2.37 | ↑ 68% |
|  | F | DopEcR | +2.36 | +2.21 | +0.15 | 7.03 | ↑ 2% |
|  |  | Dop1R1 | +0.34 | +0.45 | -0.21 | 7.03 | ↓ 3% |
|  |  | Dop1R2 | +1.34 | -1.56 | +2.90 | 7.03 | ↑ 41% |
|  |  | Dop2R | +0.79 | -2.65 | +3.44 | 7.03 | ↑ 49% |
| *αβ* | M | DopEcR | +0.55 | +3.53 | -2.98 | 4.59 | ↓ 65% |
|  |  | Dop1R1 | -0.04 | +1.61 | -1.65 | 4.59 | ↓ 36% |
|  |  | Dop1R2 | +1.53 | -0.30 | +1.83 | 4.59 | ↑ 40% |
|  |  | Dop2R | +1.74 | -0.64 | +2.38 | 4.59 | ↑ 52% |
|  | F | DopEcR | +2.09 | +2.21 | -0.12 | 7.71 | ↓ 2% |
|  |  | Dop1R1 | +1.28 | +0.45 | +0.84 | 7.71 | ↑ 11% |
|  |  | Dop1R2 | +1.87 | -1.56 | +3.43 | 7.71 | ↑ 44% |
|  |  | Dop2R | -3.13 | -2.65 | -0.48 | 7.71 | ↓ 6% |
| *α'β'* | M | DopEcR | +3.54 | +3.53 | +0.01 | 3.38 | ↑ <1% |
|  |  | Dop1R1 | -0.28 | +1.61 | -1.89 | 3.38 | ↓ 56% |
|  |  | Dop1R2 | +1.58 | -0.30 | +1.87 | 3.38 | ↑ 55% |
|  |  | Dop2R | +2.00 | -0.64 | +2.64 | 3.38 | ↑ 78% |
|  | F | DopEcR | -1.59 | +2.21 | -3.80 | 10.90 | ↓ 35% |
|  |  | Dop1R1 | -2.94 | +0.45 | -3.39 | 10.90 | ↓ 31% |
|  |  | Dop1R2 | -1.16 | -1.56 | +0.40 | 10.90 | ↑ 4% |
|  |  | Dop2R | -3.01 | -2.65 | +0.36 | 10.90 | ↑ 3% |
| *γ* | M | DopEcR | +2.13 | +3.53 | -1.40 | 2.71 | ↓ 52% |
|  |  | Dop1R1 | -0.75 | +1.61 | -2.36 | 2.71 | ↓ 87% |
|  |  | Dop1R2 | -0.38 | -0.30 | -0.08 | 2.71 | ↓ 3% |
|  |  | Dop2R | +1.15 | -0.64 | +1.78 | 2.71 | ↑ 66% |
|  | F | DopEcR | +2.74 | +2.21 | +0.53 | 7.35 | ↑ 7% |
|  |  | Dop1R1 | +1.10 | +0.45 | +0.65 | 7.35 | ↑ 9% |
|  |  | Dop1R2 | +3.53 | -1.56 | +5.09 | 7.35 | ↑ 69% |
|  |  | Dop2R | -0.29 | -2.65 | +2.37 | 7.35 | ↑ 32% |

**Supplemental Figure 1.**
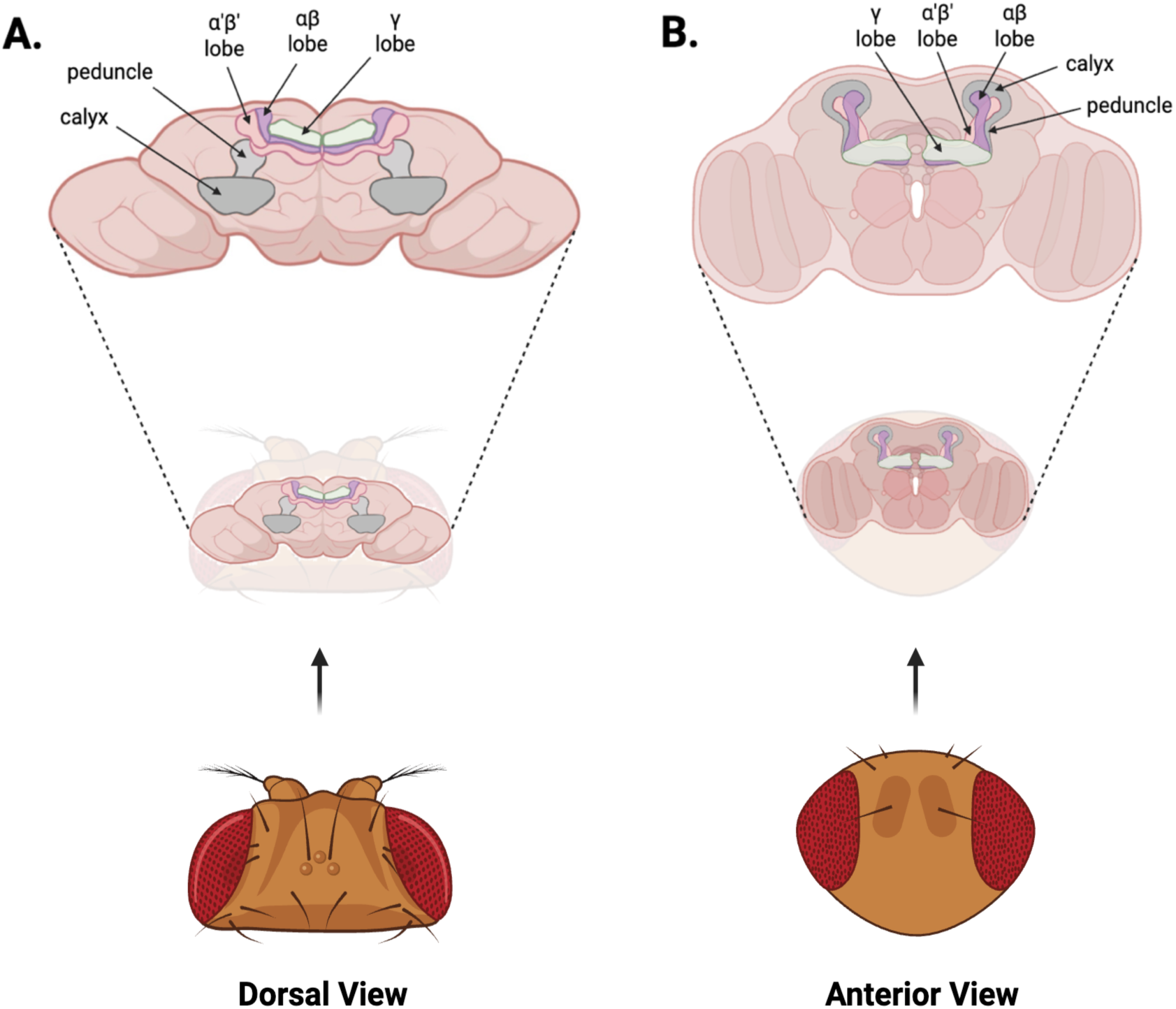
The Drosophila mushroom bodies. The mushroom body (MB) brain region is shown from dorsal **(A)** and anterior **(B)** perspectives, with major structural components labeled. Sensory information enters the MB through the calyx, which contains the dendritic arbors of intrinsic MB Kenyon cells (KCs). KC axons project anteriorly through the peduncle before branching to form the three MB lobes (αβ, α′β′, and γ). KC_αβ_ and KC_α′β′_ axons bifurcate to extend dorsally and medially, generating the L-shaped αβ and α′β′ lobes, whereas KC_γ_ axons project medially to form the horizontal γ lobes. Image created in BioRender.

**Supplemental Figure 2.**
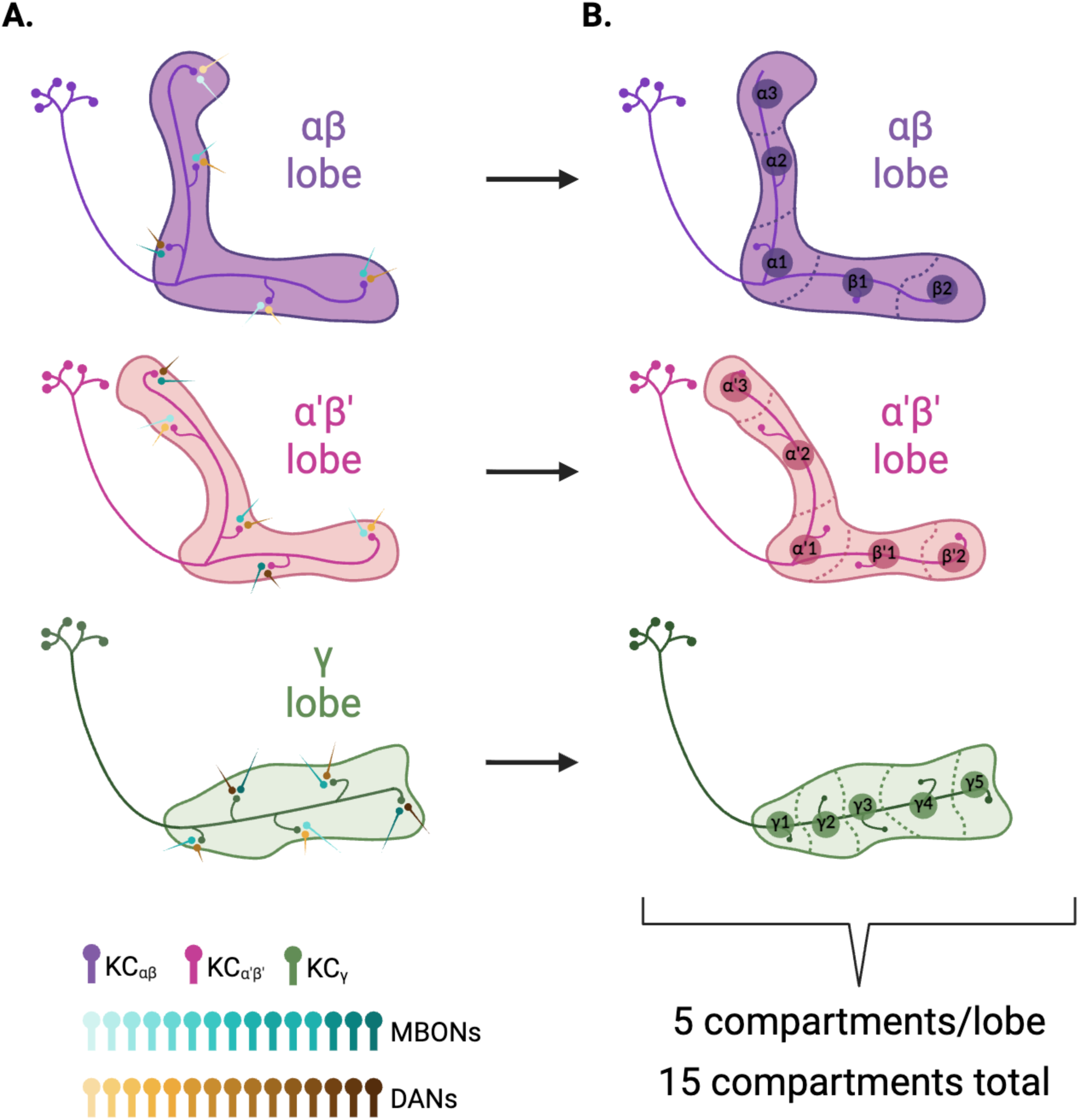
Compartmental organization of the mushroom bodies. **A.** Simplified schematics of each mushroom body (MB) lobe (αβ, α′β′, and γ), with a representative Kenyon cell (KC) shown in the corresponding colour. KC axons form *en passant* synapses with mushroom body output neurons (MBONs; teal shades) along their length, and dopaminergic neurons (DANs; amber shades) provide modulatory input at these sites. **B.** The alignment of these synaptic zones with the specialized innervation patterns of MBON and DAN subtypes defines 5 compartments per lobe, yielding a total of 15 compartments across the MB. Image created in BioRender.

**Supplemental Figure 3.**
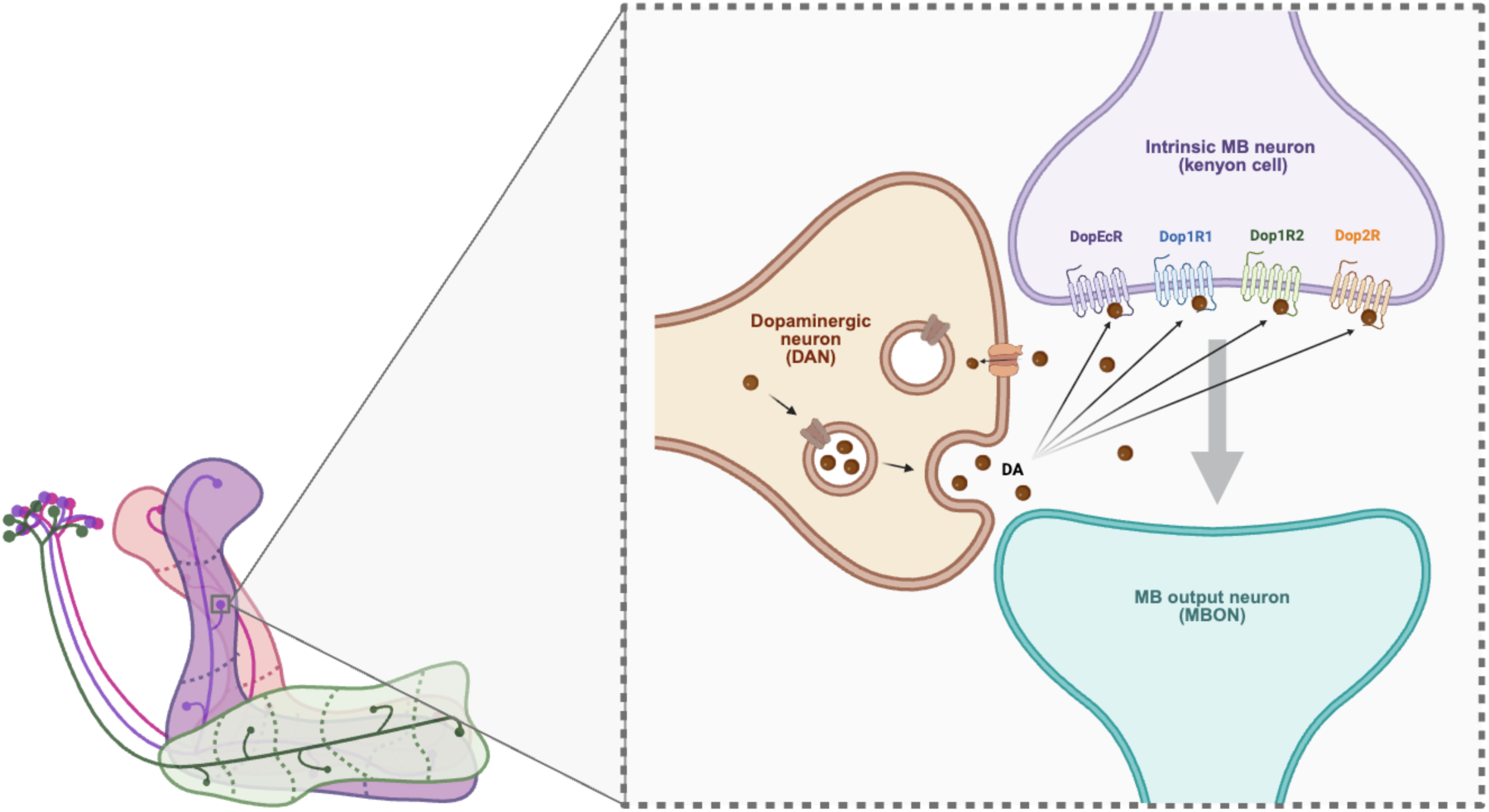
Dopaminergic modulation at KC-MBON synapses. Dopaminergic neurons (DANs) provide modulatory input at KC-MBON synapses throughout the mushroom bodies by releasing dopamine (DA; brown spheres) into the synaptic cleft to convey contextual information about prior experiences or internal state. Dopamine can bind to any of the four KC-expressed dopamine receptors (DopEcR, Dop1R1, Dop1R2, Dop2R), which are illustrated together here for simplicity but are not necessarily co-localized at the same synapse. Through these receptors, KCs integrate dopaminergic signals and adjust their presynaptic output. Such changes in KC activity alter the responses of MBONs and thus modify the information relayed to downstream circuits. Image created in BioRender.

**Supplemental Figure 4.**
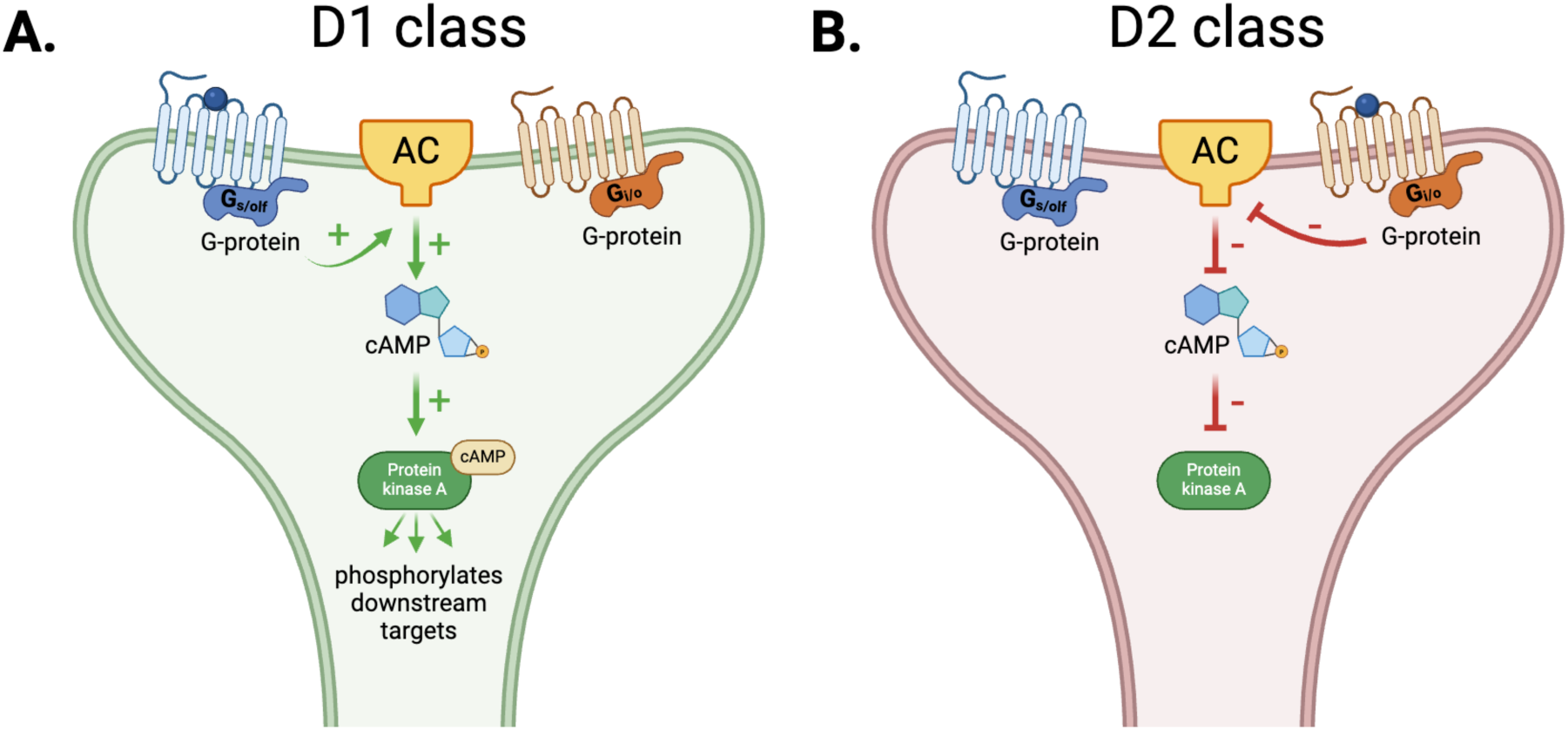
Intracellular cAMP/PKA signaling cascades initiated by dopamine binding to D1– and D2– class receptors. **A.** Signaling cascade triggered by activation of D1-class receptors. Dopamine binding leads to the activation of a stimulatory G protein, which increases adenylyl cyclase (AC) activity and elevates cyclic adenosine monophosphate (cAMP) levels. cAMP activates the catalytic subunits of Protein kinase A (PKA), resulting in phosphorylation of downstream targets and an increased likelihood of neuronal firing. **B.** Signaling cascade triggered by activation of D2-class receptors. Dopamine binding activates an inhibitory G protein that suppresses AC activity and lowers cAMP levels. Without sufficient cAMP to activate PKA, phosphorylation of downstream targets is reduced, making neuronal firing less likely. Image created in BioRender.

**Supplemental Figure 5.**
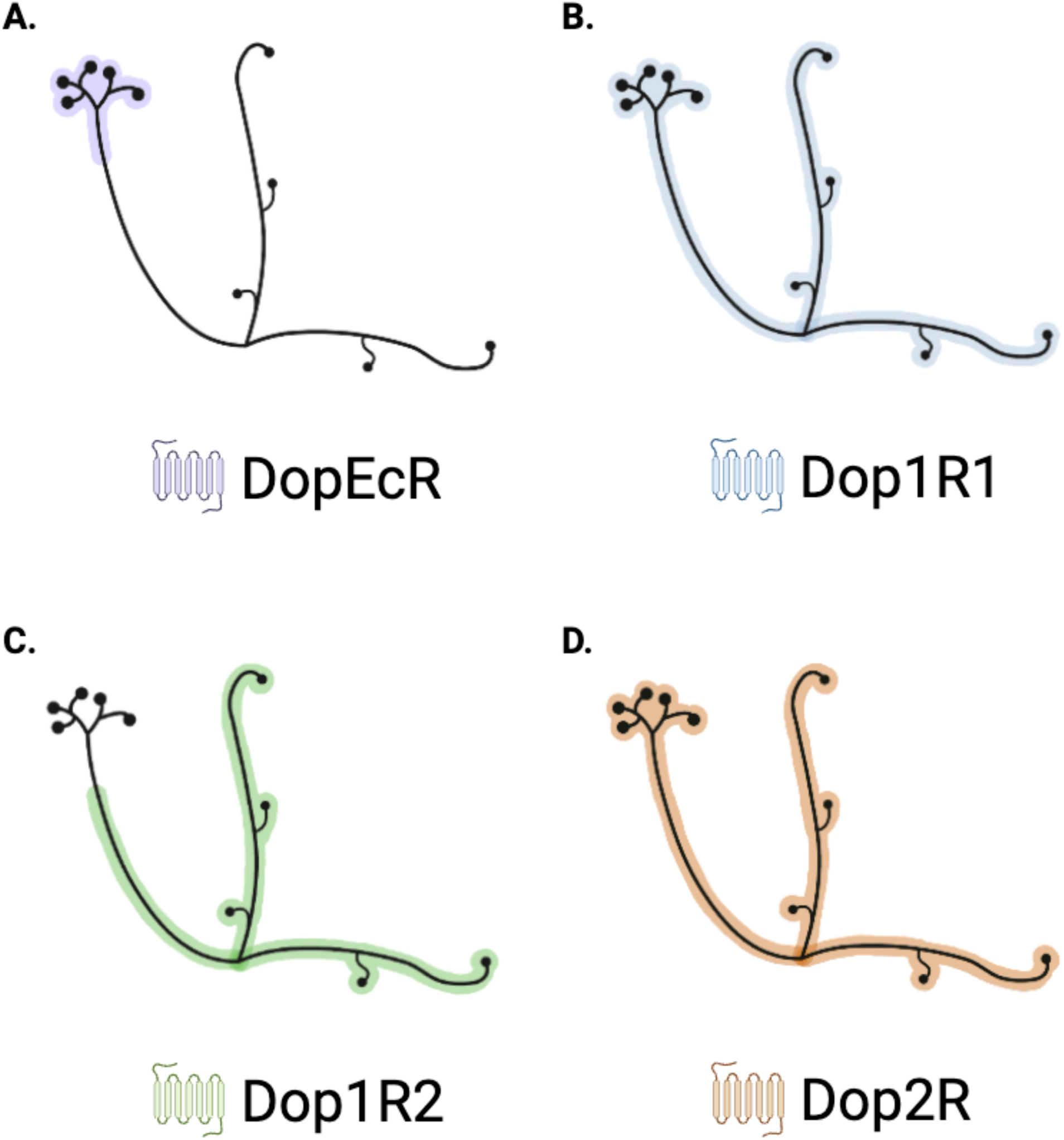
Subcellular localization of Drosophila dopamine receptors along Kenyon cell membranes in the mushroom bodies, as reported by Kondo et al. (2020). Localization patterns are illustrated using a representative αβ lobe Kenyon cell (KC). **A.** DopEcR is enriched at the axon initiation segment at the base of the peduncle. **B.** Dop1R1 displays broad distribution across the KC membrane. **C.** Dop1R2 shows a complementary pattern to DopEcR, localizing to anterior peduncular regions and distal axonal segments. **D.** Dop2R is broadly distributed along the KC membrane. Image created in BioRender.

**Supplemental Figure 6.**
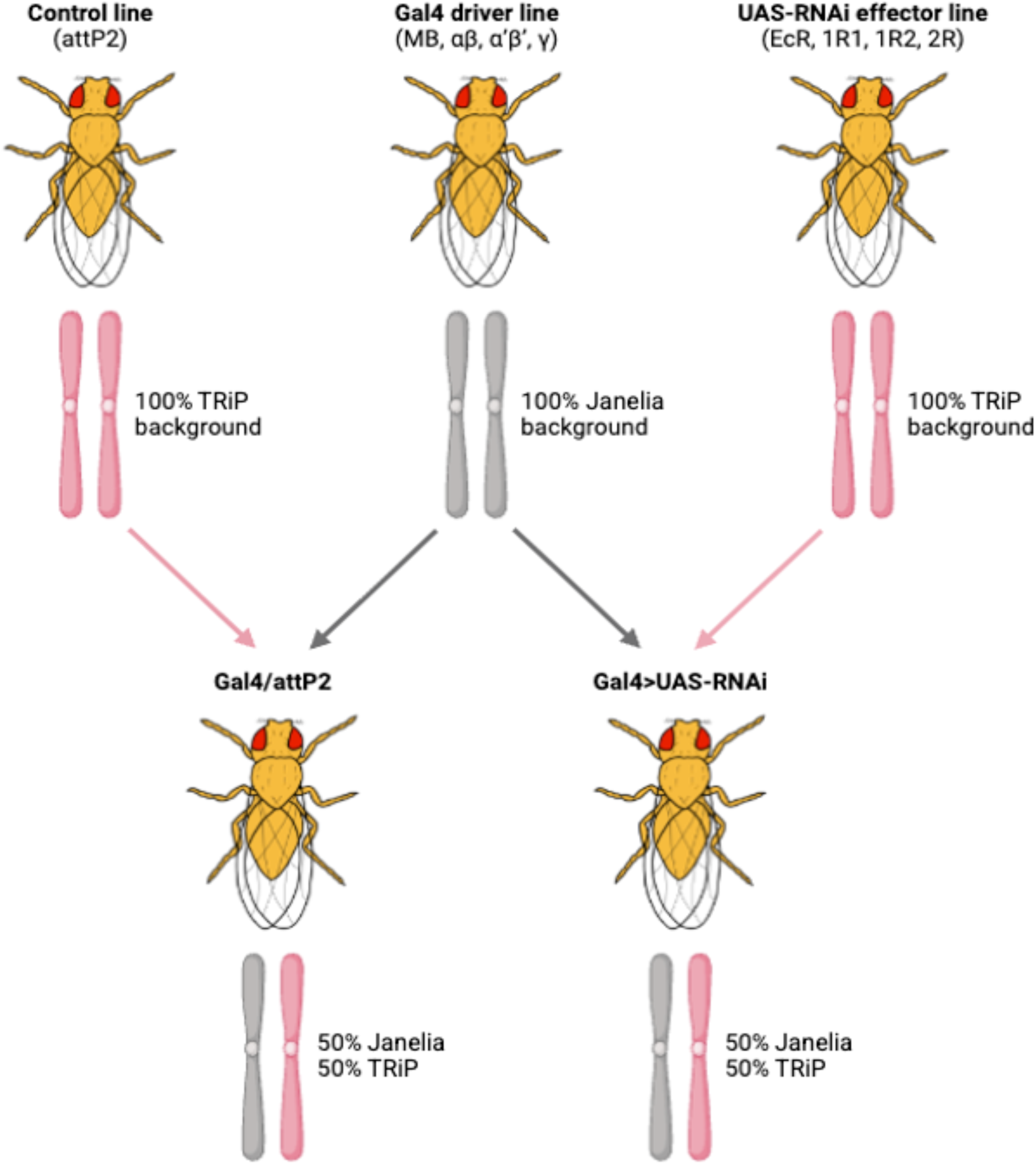
Crosses used to generate flies for behavioral assays. Parental control and UAS-RNAi lines are in a 100% TRiP genetic background. Gal4 driver lines are in a 100% Janelia background. Resulting progeny inherit a 50:50 contribution from each parental background. Image created in BioRender.

**Supplemental Figure 7.**
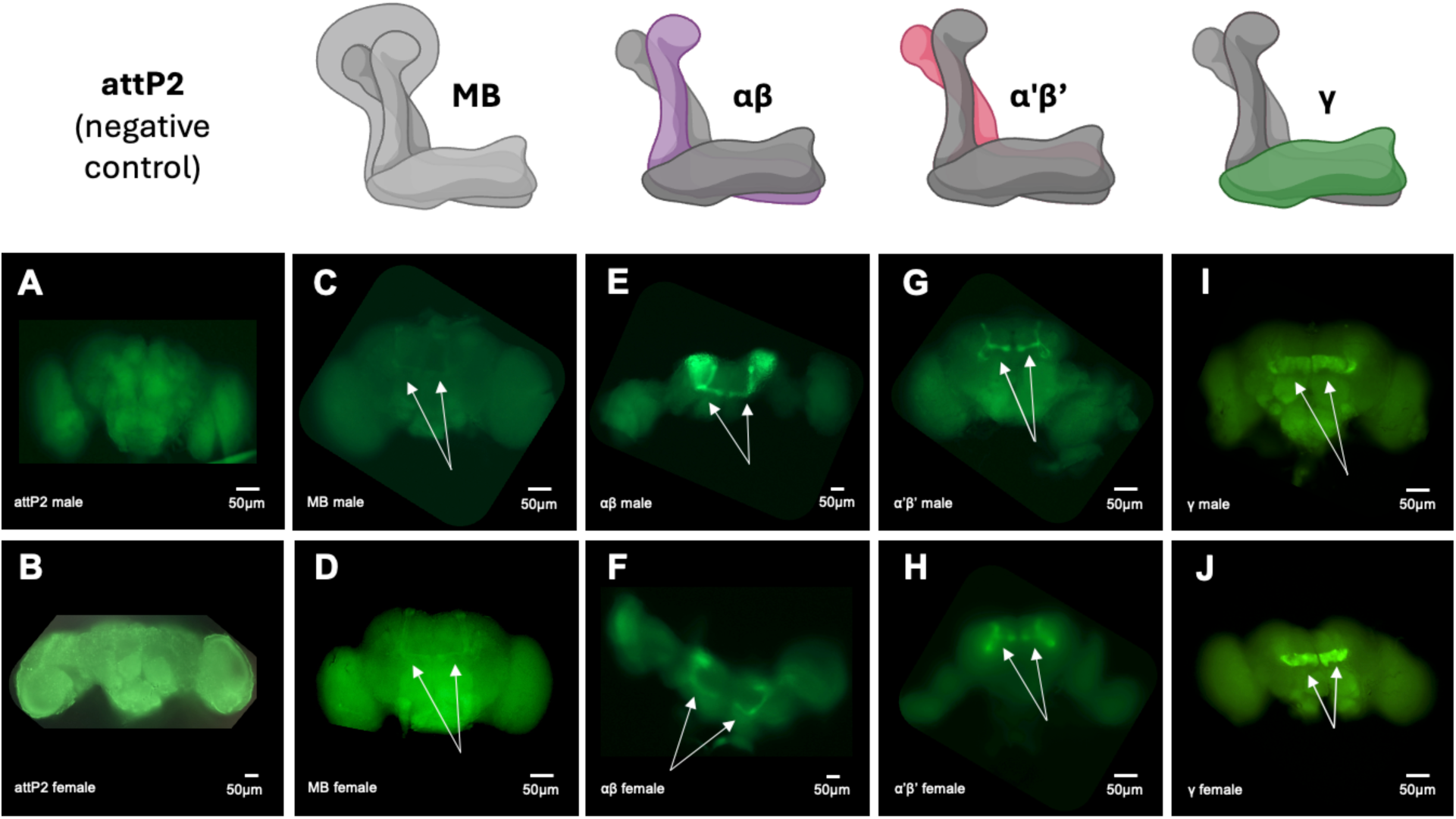
GFP expression in male and female Drosophila brains driven by mushroom body and lobe-specific Gal4 driver lines. Whole-brain fluorescence images show membrane-bound GFP expression in males (top) and females (bottom), each expressing GFP under the control of a distinct Gal4 driver. Bright green signal reflects GFP expression; relevant structures are indicated with white arrows. Autofluorescence from brain lipids accounts for diffuse background fluorescence outside the marked regions. **A-B:** *attP2* control brains with no GFP expression. Background fluorescence is present but no specific structures are visible above this signal. **C-D:** *MB* driver. **E-F:** *αβ* lobe driver. **G-H:** *α′β′* lobe driver. **I-J:** *γ* lobe driver. Schematics along the top of the image were made in BioRender.

**Supplemental Figure 8.**
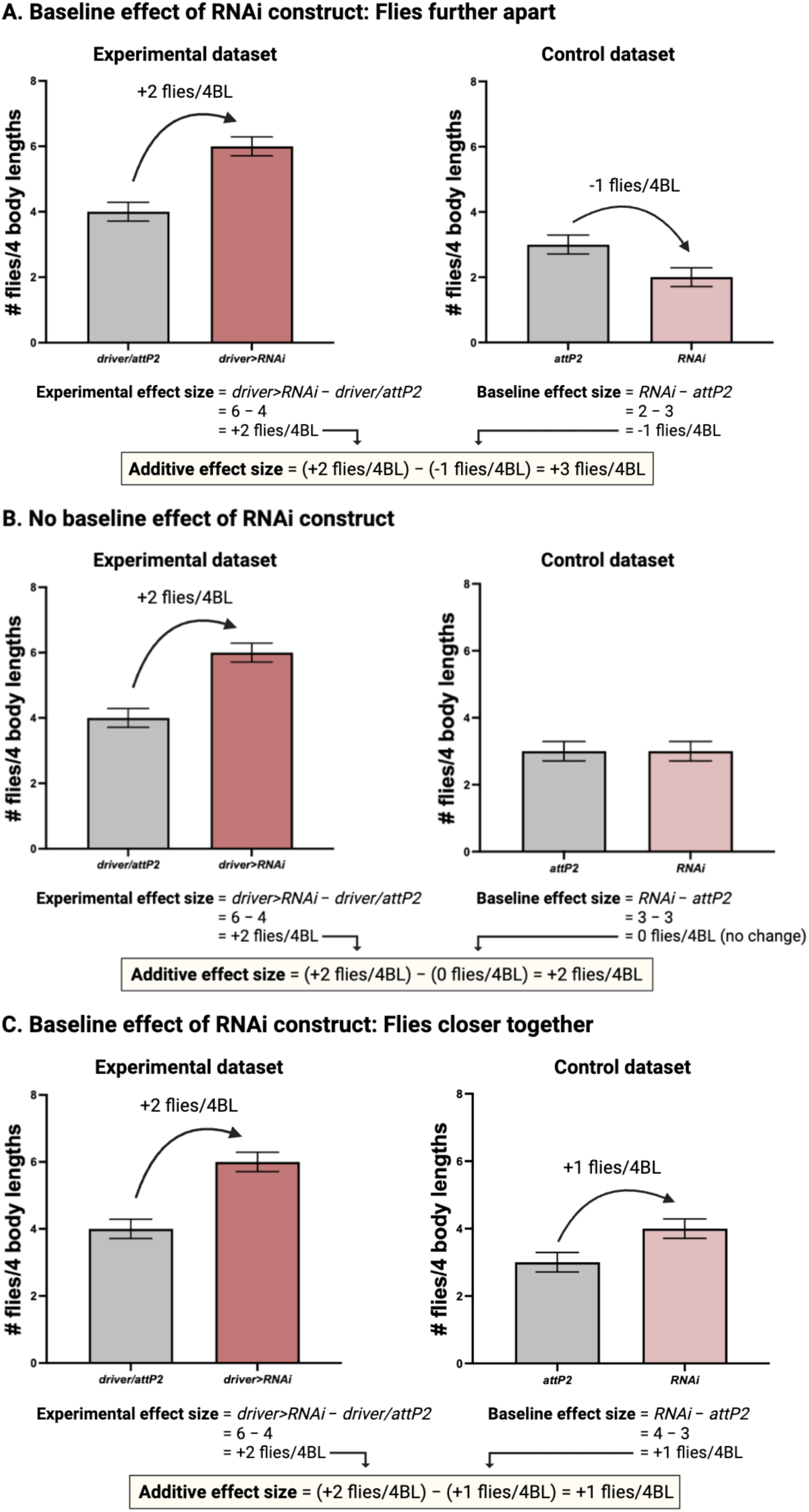
Simplified examples of effect size calculations. Three scenarios are presented to illustrate effect size quantification under different baseline conditions: **(A)** a baseline effect of the RNAi construct that increases spacing between flies, **(B)** no baseline effect of the RNAi construct, and **(C)** a baseline effect of the RNAi construct that decreases spacing between flies. Image created in BioRender.

**Supplemental Figure 9.**
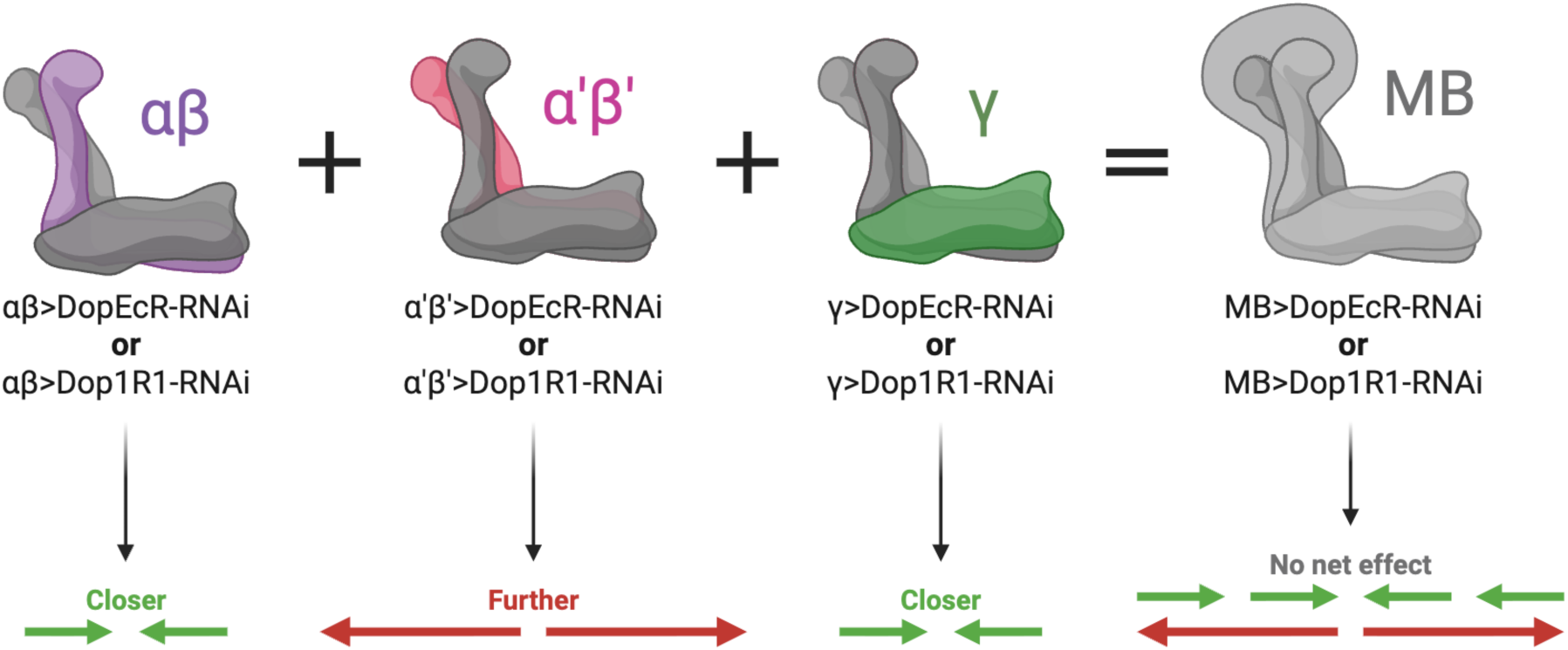
Opposing lobe-specific effects may cancel out at the MB level. Schematic illustrating the proposed mechanism by which opposing lobe-level influences of DopEcR and Dop1R1 may offset one another, resulting in no net effect when expression is manipulated across the entire MB in females. Image created in BioRender.

**Supplemental Figure 10.**
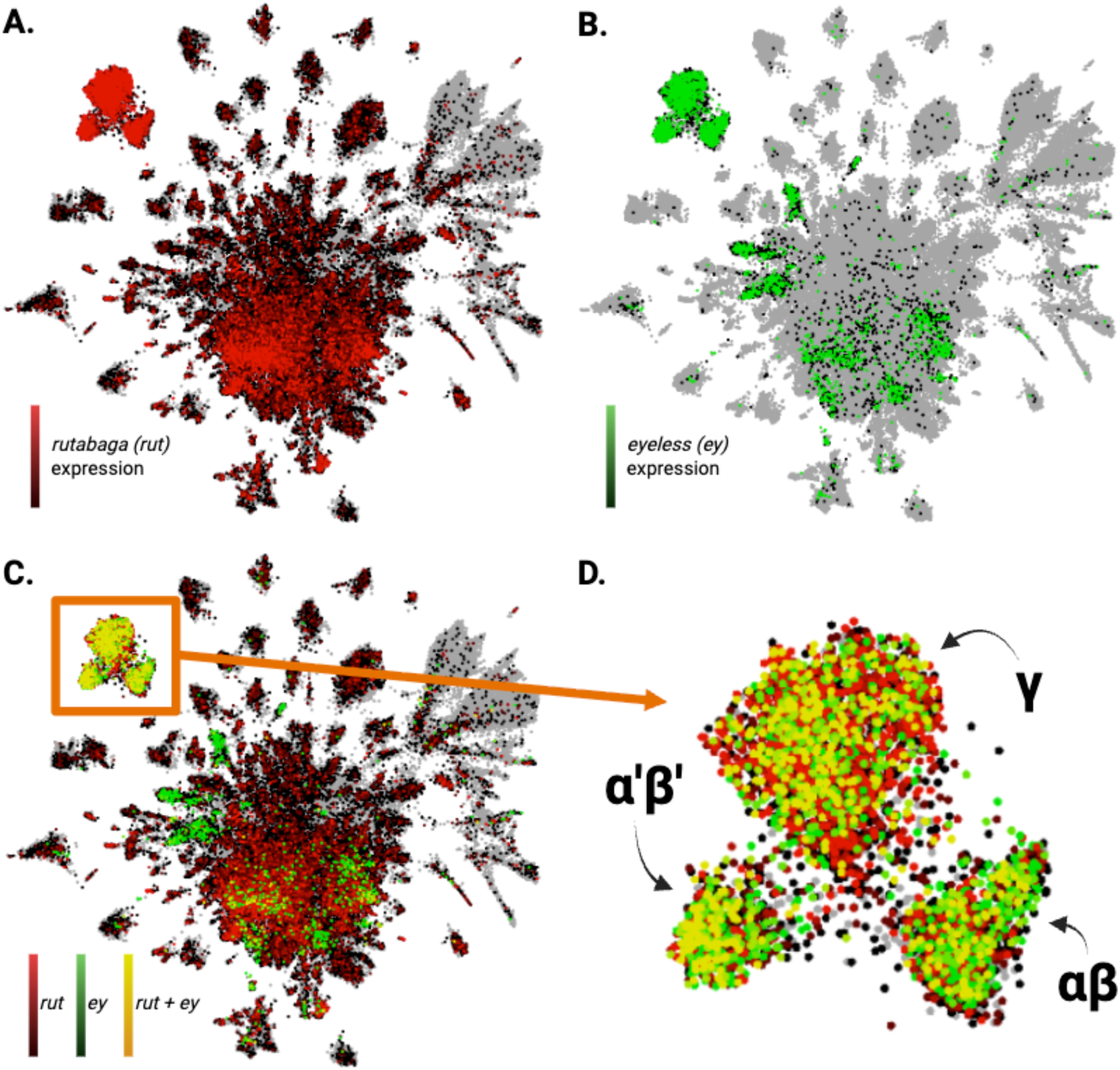
Identifying MB Kenyon cells in the FlyAtlas ScOPE UMAP plot. FlyAtlas ScOPE (Single Cell Online gene expression Profile Explorer; https://flycellatlas.org/) was used to explore gene expression patterns in Drosophila at single-cell resolution, beginning with identification of the MB Kenyon cells (Li et al., 2022). A mixed-sex adult brain dataset visualized using an HVG-based UMAP (5a3ba000_20180809_Davie_Janssens_Koldere_et al_2018_Adult_Brain.HARMONY_SCENIC) was analyzed. Expression-based plotting was used to visualize transcript levels, with higher expression represented by increased color intensity. Each point corresponds to a single cell, and clusters represent transcriptionally similar populations. Kenyon cell clusters were identified by visualizing co-expression of two established MB marker genes, *rutabaga* (*rut*) and *eyeless* (*ey*) (Han et al., 1992; Kurusu et al., 2000; Palmateer et al., 2023). **A.** *rutabaga* expression (red) is observed across much of the UMAP plot, with elevated expression in three clusters located in the upper left. **B.** *eyeless* expression (green) is more spatially restricted but similarly enhanced within the same three upper-left clusters. **C.** Overlaying *rutabaga* (red) and *eyeless* (green) expression reveals strong co-expression in these clusters, identifying them as putative MB Kenyon cells. **D.** Magnified view of the three co-expressing clusters in panel C, with tentative Kenyon cell subtype labels added based on user-suggested annotations.

**Supplemental Figure 11.**
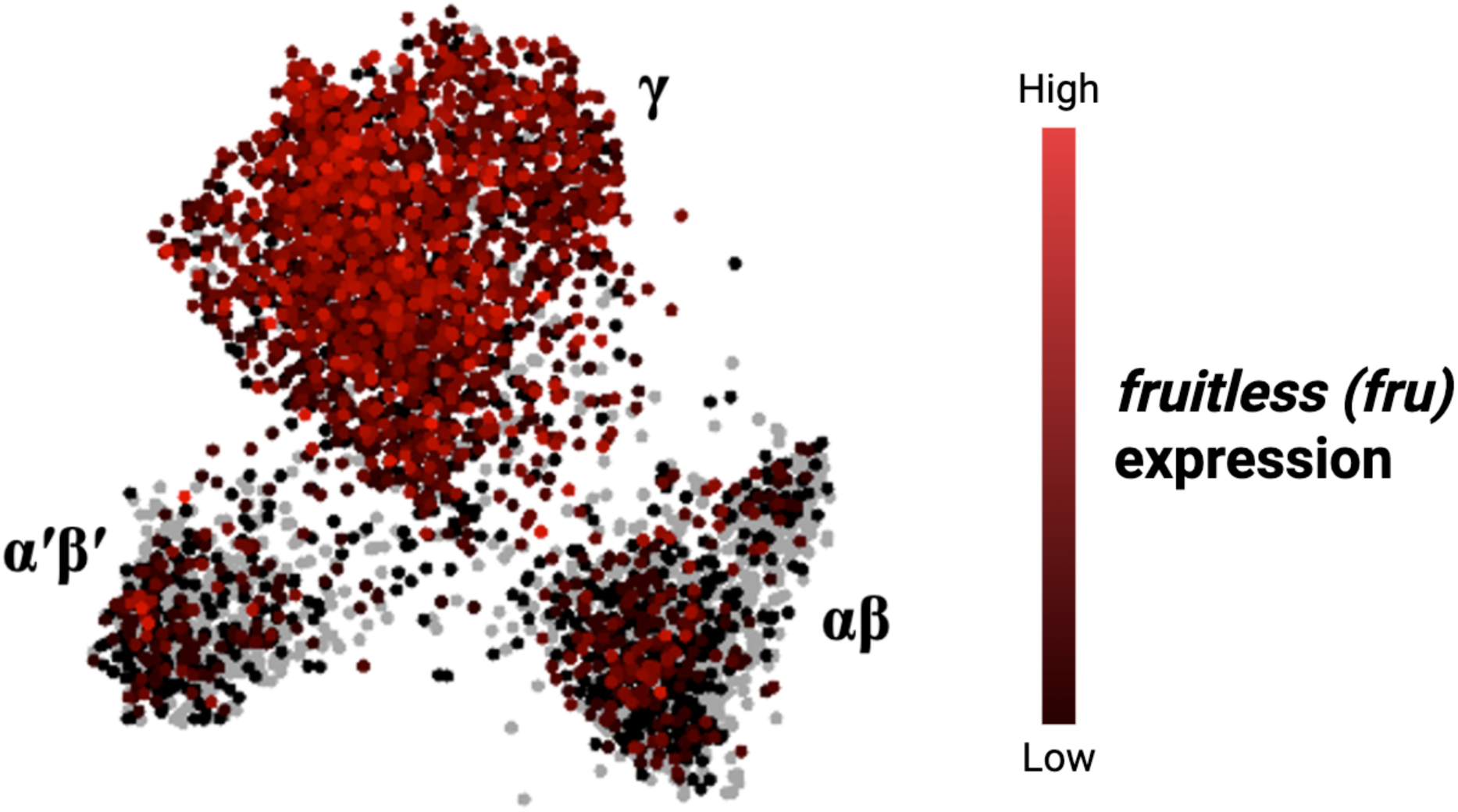
*fruitless* expression in MB Kenyon cells. Kenyon cells expressing the sex-determination factor *fruitless* (*fru*) are shown in red, with brighter colours reflecting higher expression levels. All three Kenyon cell clusters contain *fru+* neurons, with the γ cluster showing the most widespread and pronounced expression compared to αβ and α′β′ cells.

